# Repeated toxic injuries of murine liver are tolerated through microsteatosis and mild inflammation

**DOI:** 10.1101/2022.01.12.476054

**Authors:** Seddik Hammad, Christoph Ogris, Amnah Othman, Pia Erdoesi, Wolfgang Schmidt-Heck, Ina Biermayer, Barbara Helm, Yan Gao, Weronika Piorońska, Lorenza A. D’Alessandro, Fabian J. Theis, Matthias P. Ebert, Ursula Klingmüller, Jan G. Hengstler, Steven Dooley, Nikola S. Mueller

**Author notes:** Equally contributing (Seddik Hammad, Christoph Ogris, Steven Dooley, Nikola S. Mueller). **Corresponding authors, Dr. med. vet. Seddik Hammad.** Molecular Hepatology Section. Department of Medicine II. University Medical Center Mannheim. Medical Faculty Mannheim. Heidelberg University. 68167-Mannheim. Germany., **Dr. Nikola Müller.** Institute of Computational Biology. Helmholtz Zentrum München. Ingolstädter Landstr. 1. 85764-Neuherberg.

## Abstract

The liver has a remarkable capacity to regenerate and thus compensates for repeated injuries through toxic chemicals, drugs, alcohol or malnutrition for decades. However, largely unknown is how and when alterations in the liver occur due to tolerable damaging insults. To that end, we induced repeated liver injuries over ten weeks in a mouse model injecting carbon tetrachloride (CCl_4_) twice a week. We lost 10% of the study animals within the first six weeks, which was accompanied by a steady deposition of extracellular matrix (ECM) regardless of metabolic activity of the liver. From week six onwards, all mice survived, and in these mice ECM deposition was rather reduced, suggesting ECM remodeling as a liver response contributing to better coping with repeated injuries. The data of time-resolved paired transcriptome and proteome profiling of 18 mice was subjected to multi-level network inference, using Knowledge guided Multi-Omics Network inference (KiMONo), identified multi-level key markers exclusively associated with the injury-tolerant liver response. Interestingly, pathways of cancer and inflammation were lighting up and were validated using independent data sets compiled of 1034 samples from publicly available human cohorts. A yet undescribed link to lipid metabolism in this damage-tolerant phase was identified. Immunostaining revealed an unexpected accumulation of small lipid droplets (microvesicular steatosis) in parallel to a recovery of catabolic processes of the liver to pre-injury levels. Further, mild inflammation was experimentally validated. Taken together, we identified week six as a critical time point to switch the liver response program from an acute response that fosters ECM accumulation to a tolerant “survival” phase with pronounced deposition of small lipid droplets in hepatocytes potentially protecting against the repetitive injury with toxic chemicals. Our data suggest that microsteatosis formation plus a mild inflammatory state represent biomarkers and probably functional liver requirements to resist chronic damage.

**Graphical abstract:** 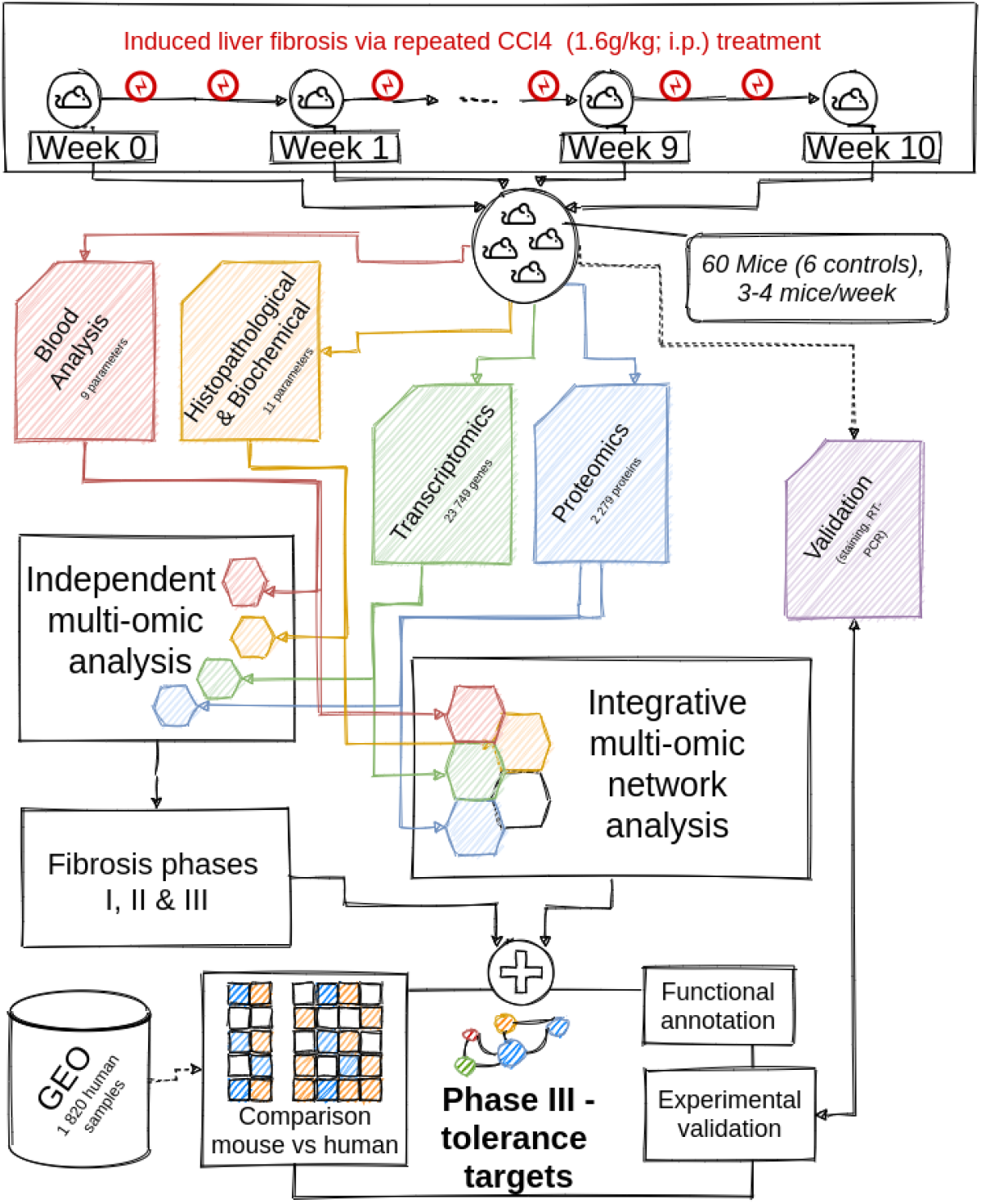

The datasets generated via transcriptomics, proteomics as well as blood, histopathological and biochemical analysis were analyzed in an independent and integrative manner. The independent analysis was performed via state-of-the-art statistical approaches i.e. differential and consistently regulated genes and proteins. Combining the results identified three fibrosis phases. Using the KiMONo algorithm, a fibrosis specific multi-omic network was inferred. Within this network we identified several nodes connecting phase III specific features forming 13 distinct multi-omic modules suggesting a tolerance scheme. Some of these modules were experimentally validated and compared to 11 independent human studies of various liver diseases.

## INTRODUCTION

Independent of etiology, chronic liver injury results in constant necrosis and apoptosis of liver cells. This triggers a process characterized by infiltration and activation of the immune cells in the liver leading to both inflammatory and wound healing responses as well as fibrogenesis (Bataller and Brenner, 2005; Seki and Schwabe, 2015). Repeated injuries of the liver tissue cause a gradual replacement of the normal liver with fibrotic tissue and lead, in a long-term process, to cirrhosis (Pellicoro et al. 2015; Zhou et al. 2014). In addition, hepatic stellate cells (HSC) change to a myofibroblast phenotype characterized by high proliferative activity, expression of ECM components, gain of contractility, chemotaxis, and migratory properties, and the production large amounts of growth factors and profibrogenic cytokines promoting fibrogenesis i.e. TGF-β (Friedmann, 2008). Maintaining quiescence and activation of HSC is a highly dynamic and complex process (Krizhanovsky et al. 2008; Kisseleva et al. 2012; Mercado-Gómez et al. 2020). Accordingly, the progression and regression of liver disease in response to repetitive injuries at the patient’s level are also highly variable (Bedossa et al. 2003; Sun et al. 2020). Fortunately, as shown previously in experimental and clinical settings (Sohrabpour et al. 2012), liver fibrosis and even the early stages of cirrhosis are reversible. Clinically, these liver diseases develop in a long-term process, taking upto 50 years (Pellicoro et al. 2015; Zhou et al. 2014). During this period, the liver can cope with many repeated small injuries induced by toxic chemicals, drugs, alcohol or a high fat diet. Therefore, it is crucial to gain deeper insights into differences between acute and chronic liver response programs that show avenues of potential targets as well as a time window for therapeutic intervention.

Up to date, studies on liver diseases in response to injury primarily compared stable disease states, like resistant versus susceptible mouse strains (Tuominen et al. 2021). Tuominen and co-workers examined by transcriptomics the susceptibility of 98 mouse strains (693 livers) to 6 weeks of CCl_4_ administration (Tuominen et al. 2021). In this report, the top 300 down-regulated targets were assigned to metabolic pathways i.e. biological oxidations, steroids and lipids (Tuominen et al. 2021). In addition, several studies analyzed the responses of the liver to CCl_4_ administration for 4-6 weeks of administration (Krizhanovsky et al. 2008; Meng et al. 2012; Cubero et al. 2016; Liu et al. 2020). However, dynamic changes in the responses of the liver upon repeated injuries have previously not been addressed.

To study complex molecular processes occurring at the cellular and tissue levels, high-throughput omics technologies provide an unbiased view. Typically, one omic level of choice is used, and state-of-the-art data analysis strategies are applied. Using only a single level of measurement, e.g., transcriptomics or histopathology, limits the detection of possible alterations to one type, and thus captures changes only for a small subset of possible components interacting and steering the disease progression (Ogris et al. 2021). To obtain a holistic view and reduce the single-level technical effects, studies utilizing multiple omic measurements are employed to support findings or identify targetable molecules in a specific disease context (Mardinoglu et al. 2018; Mercado-Gómez et al. 2020). Approaches using multi-level analysis outperform single-level analysis, as those are limited in their information content to one level and therefore may lead to missing and misinterpreted results (Domenico et al. 2015, Schmitt et al. 2013). Recently, we developed an integrative approach that is able to exploit the complexity of ‘cross-omics’ relationships of multi-level data, called KiMONo (Ogris et al. 2021). The method accounts for the limitations of the level-wise analysis by inferring condition-specific multi-omic networks. By explicitly associating, for example, a gene’s expression to its coded protein expression in addition to the expression of protein interaction partners, the resulting network captures the regulatory footprint that occurs on multiple levels. The created networks consist of genes, proteins, or blood parameters represented as nodes, linked if they are affected similarly by the underlying condition.

In this study, we delineated three phases of liver responses, namely, initiation, progression, and tolerance to repetitive administration of a toxic chemical by utilizing existing state-of-the-art multi-omic data analysis methods. To that end, we generated time-resolved measurements from blood and livers of CCl_4_-exposed mice over a time course of 10 weeks and analyzed the dynamics of liver responses at the transcriptomic, and proteomic levels, including blood parameters as indicators for organ functionality and phenotypic information (Figure 1a). To investigate this large multi-level dataset, we used a level-wise and an integrative analysis strategy employing KiMONo. Based on this analysis, we identified 13 functional modules, i.e. lipid metabolism regulated by differentially expressed genes and proteins during the tolerance phase fostering the protective response and thus survival of the animals. Based on KiMONo’s predictions, we found that the liver response of lipid accumulation was associated with tolerance. Collectively, several tolerance-related targets were analyzed in publically available human datasets and consistent deregulation was observed. Besides understanding the dynamics of liver fibrosis, these pathways were identified and validated for new potential therapeutic targets of liver fibrosis.

**Figure 1:**
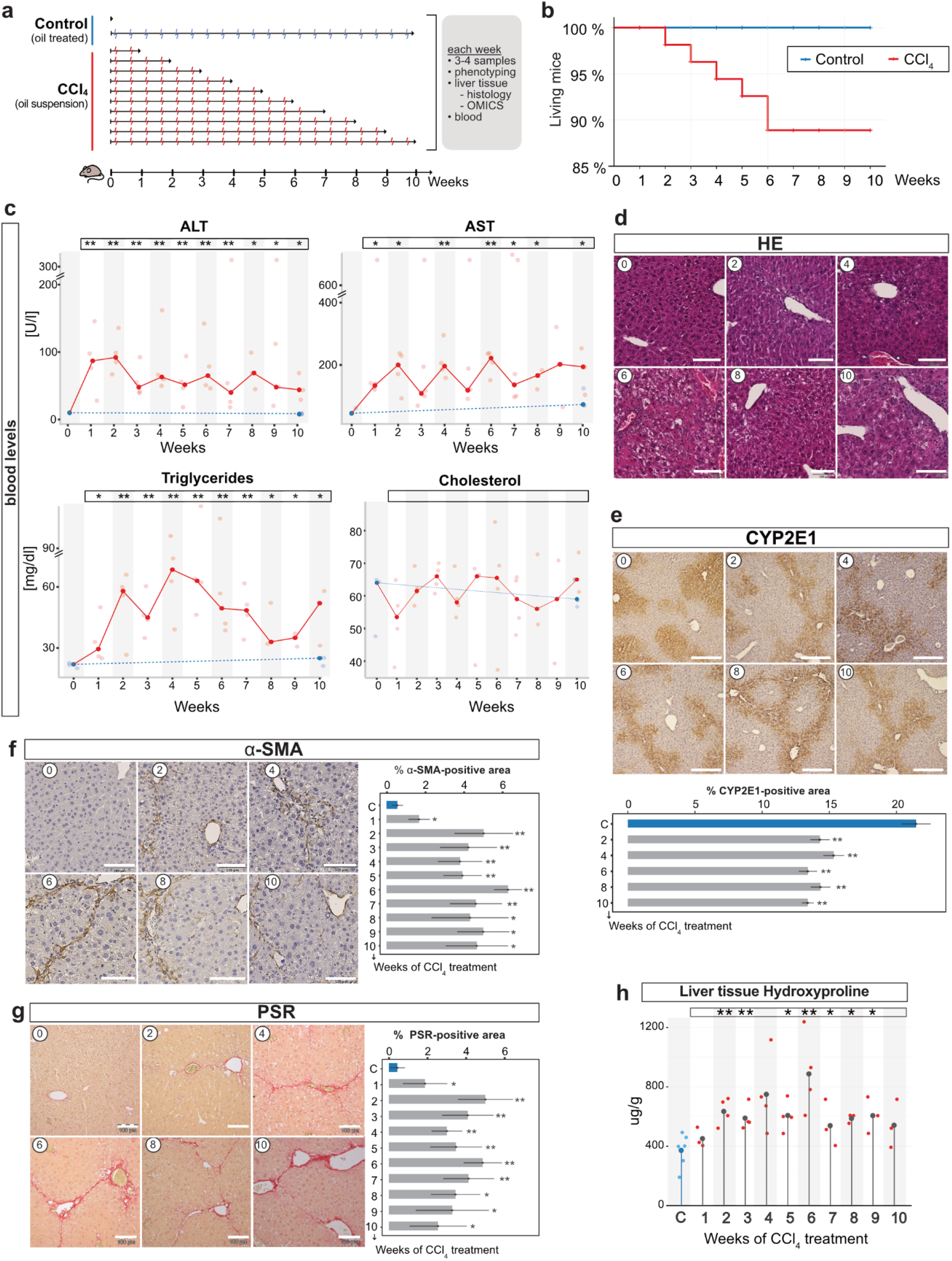
Blood, histopathological and biochemical analysis of liver fibrosis dynamics upon CCl_4_ injections. (a) Induction of experimental liver fibrosis by CCl_4_ in mice twice per week for 10 weeks. Blood and liver were collected every week for further multi-levels analysis. (a) Kaplan-Meier curve for survival analysis. (c) Blood based analysis of ALT, AST, TGs and Chol in a time-resolved manner. Liver sections were prepared from liver tissue samples of control and mice with CCl_4_ treatment for 10 weeks and HE (d), CYP2E1 (e), α-Sma (f) and PSR (g) staining was performed and positive signals were quantified as a percentage of total area. (h) Biochemical analysis of HYP level in a time-resolved manner. Results were expressed as mean of 3-6 mice± SD, and were compared by two-way ANOVA test. *p <0.05, **p <0.01 compared to 0 week (control). CCl_4_, carbon tetrachloride; ALT, alanine aminotransferase; AST, aspartate aminotransferase; Chol, cholesterol; Cyp2E1, Cytochrome P4502e1; TG, triglycerides; HE, hematoxylin & eosin; α-Sma, alpha-smooth muscle actin; PSR, picrosirius red; HYP, hydroxyproline. n=3-6 per group.

## MATERIALS AND METHODS

### Animal models of hepatic fibrosis

Adult male C57Bl/6N mice were obtained from the Janvier labs (France), housed three per cage in a temperature-controlled (24°C) room with a 12-h light/dark cycle, and given *ad libitum* access to water and laboratory diet (Ssniff, Germany). Mice were maintained for 7 days before carbon tetrachloride (CCl_4_) intoxication. The dose of CCl_4_ (Sigma-Aldrich, Cat. no. 319961) was 1.6 g/kg body weight (Hammad et al. 2017) and was prepared as follows: to 3 ml of olive oil, 1 ml of CCl_4_ was added and mixed well. Mice received CCl_4_ intraperitoneally twice per week. In a time-resolved manner including 1, 2, 3, 4, 5, 6, 7, 8, 9 and 10 weeks, mice were sacrificed at day two after the last CCl_4_ injection. Identical concentration of olive oil was injected for control groups for 10 weeks (Figure 1a). At the indicated time point, blood and livers were harvested. The liver lobes were separated as follows: the caudate lobe (for hydroxyproline), right lobe (for proteomics and transcriptomics) and median lobe (cryosectioning) were freshly frozen in liquid nitrogen and stored at −80°C. Further, left liver lobes were fixed in 4% paraformaldehyde (PFA) and embedded in paraffin for histopathological investigations. The experimental protocols with animals were carried out in full compliance with the guidelines for animal care and were approved by the Animal Care Committee from the German government (Animal permission number: 35.9185.81/G-216/16).

### Clinical Chemistry

Blood was collected in Li-Heparin vials from the retrobulbar plexus and centrifuged at 13,000 rpm at 4°C for 6 min. Plasma was subsequently stored at − 80°C until further analysis. Then, alanine aminotransferase (ALT), aspartate aminotransferase (AST), alkaline phosphatase (ALP), glucose (Gluc), triglycerides (TG), total protein (Prot), blood urea nitrogen (BUN) and cholesterol (Chol) were measured using a Hitachi automatic analyzer (Core facility-Medical Faculty Mannheim, Germany).

### RNA isolation and transcriptome analysis

Pieces of the right liver lobes were used to perform mRNA isolation with the InviTrap®Spin Universal RNA Mini Kit from Stratec (1060100300, Birkenfeld, Germany) according to the manufacturer. RNA concentration and integrity were summarized in Supporting table 1. Transcriptomics using the isolated mRNA from liver tissues (0, 2, 4, 6, 8 and 10 weeks; n=3 per time point) was performed by Affymetrix GeneChip®Mouse Gene 2.0 ST Arrays (902118). Affymetrix-based transcriptomics was performed at Core Facility-University Clinic Mannheim (Germany; http://zmfsrv1.medma.uni-heidelberg.de/apps/zmf/Affymetrix/).

### Transcriptomic data preprocessing and bioinformatic analysis

Gene expression data obtained by the whole-transcript array GeneChip Mouse Gene 2.0 ST were pre-processed using ‘affyPLM’ packages of the Bioconductor Software (Huber et al. 2015). Genes with the strongest evidence of differential expression were obtained using a linear model provided by the limma-package (Ritchie et al. 2015). Data obtained from untreated mice were used as a reference. To annotate the microarrays, a custom chip definition file version 22 from Brainarray (Dai et al. 2005) based on Entrez ID’s was used. A false-positive rate of α = 0.05 with false discovery rate (FDR) correction and a fold change greater than 1.5 was taken as the level of significance. To unravel patterns in the gene expression data for different pathways heatmaps the ‘ComplexHeatmap’ (Gu et al. 2016) package was used.

### Reverse transcription polymerase chain reaction (RT-PCR)

Using RNA isolated for transcriptomics analysis, cDNA was produced from oligo(dT)18 primer (SO132, Thermo Scientific, Massachusetts, USA), dNTP Mix (R0191, Thermo Scientific Massachusetts, USA), and RevertAid H Minus Reverse Transcriptase (EP0451, Thermo Scientific, Massachusetts, USA), and then used for real-time (rt)-PCR (5× HOT FIREPol EvaGreen qPCR Mix Plus (ROX), 08-24-00020, Solis BioDyne, Tartu, Estonia) in a StepOne machine. Sequences of primer pairs were listed in Supporting table 2. All primers were purchased from Eurofins Genomics (Ebersberg, Germany). The mRNA expression levels of the detected genes were normalized to that of *Ppia*.

### Sample preparation for proteome analysis

The proteome profiling of liver tissue was performed on selected time points (0, 1, 2, 4, 6, 8, and 10 weeks; n=3 per time point). Liver tissue was powdered using a Micro-Dismembrator (B.Braun, Micro-Dismembrator U Ball Mill), and approximately 10 mg aliquots of tissue powder were lysed in 100 μl SDS buffer (4% SDS, 1x Halt protein inhibitor, 40 mM TCEP, 160 mM CAA, 200 mM TEAB) by sonication on ice (60 sec, 80% amplitude, 0.1 sec off/0.5 sec on) and centrifugation (15 min, 14,000 rpm, 4°C). The supernatant was incubated at 95°C for 5 min and 70°C for 30 min for reduction and alkylation. Protein concentrations were determined by BCA assay, and 20 μg of total protein was used for subsequent tryptic digestion. Samples were prepared using a modified version of the Single-pot, solid-phase-enhanced sample preparation (SP3) protocol (Hughes et al. 2014). Briefly, a mix of Sera-Mag SP3 beads was added to the protein samples in a 10:1 SP3 beads/protein (wt/wt) ratio. Acetonitrile was added for a final concentration of 70% organic, and the mix was incubated for 18 min at room temperature (RT). Protein-bound beads were isolated on a magnetic rack and washed twice with 70% ethanol. A third wash was performed using 100% acetonitrile (ACN). Beads were air-dried and reconstituted in 100 mM TEAB buffer containing Trypsin Gold (Promega) in a 1:25 enzyme/protein (wt/wt) ratio. Protein digestion was performed for 14 h at 37°C. The digested peptides were dried by vacuum centrifugation and stored at −20°C until further use. A sample pool, consisting of 5 μg of each sample in the dataset, was generated. Peptides were then reconstituted in 50 mm HEPES (pH 8.5), and TMT10-plex reagents (ThermoFisher) were added to the samples (stocks dissolved in 100% ACN) in a 1:10 sample/TMT (wt/wt) ratio. The peptide–TMT mixture was incubated for 1 h at RT, and the labeling reaction was stopped by addition of 5% hydroxylamine to a final concentration of 0.4%. Different samples were combined and the TMT-plexes were fractionated into 6 fractions using stage-tip Strong cation-exchange (SCX) fractionation. Stage-tips were manually prepared using 3 discs of SCX resin (Empore) and conditioned with MeOH, followed by 80% ACN, 0.5% AcOH; 0.5% AcOH; 500 mM NH4AcOH, 0.5% AcOH, 30% ACN; 20mM NH4AcOH, 0.5% AcOH, 30% ACN, and was equilibrated with 0.5% AcOH successively. Before SCX fractionation, samples were reconstituted in 0.5% AcOH, sonicated, incubated on a shaker, and loaded onto the stage tips by centrifugation. Loaded stage-tips were washed with 0.5% AcOH. For elution, descending concentrations of elution buffers (20/40/70/100/250/500 mM of NH4AcOH, 0.5% AcOH, 30% ACN) were used and the flow-through was collected. Fractions were dried by vacuum centrifugation and stored at −20°C

### LC-MS/MS Measurements

Nano-flow LC-MS/MS was performed by coupling an EASY-nLC (Thermo Scientific, USA) to a Q Exactive HF-X – Orbitrap mass spectrometer (Thermo Scientific, Germany). The fractions were dissolved in 11 μl loading buffer (0.1% formic acid, 2% ACN in LC-MS grade water), sonicated and incubated on a shaker. 5 μl of each fraction was used for each measurement. Peptides were delivered to an analytical column (75 μm × 30 cm, packed in-house with Reprosil-Pur 120 C18-AQ, 1, 9 μm resin, Dr. Maisch, Ammerbuch, Germany) at a flow rate of 3 μl/min in 100% buffer A (0.1% formic acid in LC-MS grade water). After loading, peptides were separated using a 120 min stepped gradient from 6% to 50% of solvent B (0.1% formic acid, 80% ACN in LC-MS grade water; solvent A: 0.1% formic acid in LC-MS grade water) at 350 nL/min flow rate. The Q Exactive HF-X was operated in data-dependent mode (DDA), automatically switching between MS and MS2. Full-scan MS spectra were acquired in the Orbitrap at 120,000 (m/z 200) resolution after accumulation to a target value of 3,000,000. Tandem mass spectra were generated for up to 18 peptide precursors in the Orbitrap (isolation window 0.8 m/z) for fragmentation using higher energy collisional dissociation (HCD) at normalized collision energy of 32% and a resolution of 45,000 with a target value of 50,000 charges after accumulation for a maximum of 96 ms.

### Protein identification, quantification and statistical analysis

Raw MS spectra were processed by MaxQuant (version 1.6.0.1) for peak detection and quantification. MS/MS spectra were searched against the Uniprot *mus musculus* reference proteome database (downloaded on October 22nd, 2018) by Andromeda search engine enabling contaminants and the reversed versions of all sequences with the following search parameters: Carbamidomethylation of cysteine residues as fixed modification and Acetyl (Protein N-term), Oxidation (M) as variable modifications. Trypsin/P was specified as the proteolytic enzyme with up to 3 missed cleavages allowed. The mass accuracy of the precursor ions was decided by the time-dependent recalibration algorithm of MaxQuant. The maximum false discovery rate (FDR) for proteins and peptides was α = 0.01 and a minimum peptide length of eight amino acids was required. Quantification mode with isobaric labels (TMT 10plex) was selected. All other parameters are defined as default settings in MaxQuant.

### Proteomic data preprocessing and bioinformatic analysis

By filtering out contaminant proteins, as indicated by MaxQuant analysis, and using the corrected reporter intensity for analysis, all proteins with at least one missing expression value in one of the samples were removed, leaving 2278 proteins for subsequent analysis. This stringent cutoff was chosen as visual inspection of the data showed that missingness increases with reporter intensity, thus protein expression. We computed the ratio per *p* protein *j* on the basis of its sample’s expression *s* (pj,s) and the samples-matching protein expression in the TMT10plex isotope reference channel *r* (rj) according to (1+pj,s)/(1+rj). One outlier sample was removed after this stage. Robust quantile normalization was applied to the data using the MSnSet R package, resulting in final normalized expression ratios used for statistical analysis. For identification of differentially regulated proteins per time point (1, 2, 4, 6, 8 and 10 weeks) when compared to the control time point (0 week), we used a multivariate linear model with time-point specific dummy variables and protein ratio as a response. To call differential expression, we subjected p-values of all proteins separately per coefficient to multiple testing correction with the Benjamini Hochberg procedure, also referred to as False-Discovery Rate. Differential expression per time-point was set to FDR below α = 0.05.

### Multi-omics network inference and analysis

The multi-omic network inference was performed using KiMONo (Ogris et al 2021). This novel versatile tool can use any kind and any amount of omic data by leveraging prior knowledge. By doing so KiMONo generates a multi-level network around an omic type of interest, simplifying downstream analysis, i.e. pathway analysis. In the final multi-level network nodes represent features like proteins, genes or clinical variables and the connections between them denote effects identified within the input data. Here, we used KiMONo to generate three networks centered around the three given data types and combined them to enhance the signal within the time-resolved data. This was done by only reporting effects that were found in all three multi-omic networks. To infer each network, we used three different priors providing information about already known relations between the transcriptomic, proteomic, and clinical data. In KiMONo the priors serve as a rough blueprint, reducing the complexity and improving the algorithm’s performance. The first priority was obtained via the Biomart tool annotating genes to proteins. The second priority is based on the BioGRID database to include information about protein-protein relations (Oughtred et al. 2019). As a third priority, we used all previous annotations to identify indirect gene-protein interactions. This was done by using BioGRID interactions information to also annotate genes to proteins of their coding protein. Finally, we set off to generate a prior which would annotate the omic information to our clinical data. Therefore, we first inferred the transcriptomic and proteomic centered networks and used the information about all potential effects of clinical features as prior for the clinical centered network inference. The importance of network nodes was estimated via the network’s betweenness centrality. This measure is estimated by determining the shortest path between all nodes within the graph. The betweenness centrality for a node is then estimated by the number of shortest paths that pass through a node. Functional annotation of network modules was performed by using the online network-based pathway enrichment tool PathwaX (Ogris et al. 2016) and the KEGG (Kanehisa et al. 2016) and Reactome (Jassal et al. 2019) pathways. Significantly enriched pathways (FDR < 0.05) were manually curated to identify overarching functional themes which were assigned as labels to modules. Moreover we also manually annotated small modules (2-3 elements) via literature search.

### Human ortholog comparison

We obtained 1034 human samples across 11 liver disease related datasets via NCBI’s Gene Expression Omnibus (GEO), see Supporting table 3. Significant differentially expressed (DE) genes (FDR < 0.1) have been extracted by using the GEO2R interface and its default settings. Further, we identified mouse orthologous genes using the Inparanoid 8 database (Sonnhammer and Östlund 2015).

### Hepatic hydroxyproline determination

Hydroxyproline (HYP) was determined colorimetrically in triplicates from snap-frozen liver lobes as described in Fels (1958) with modifications. Briefly, approximately 100 mg of tissue from the caudate liver lobe was homogenized and hydrolyzed in 2 ml of 6 N HCl at 110°C for 16h. HYP content was then measured photometrically at 558 nm. Based on relative HYP (per 100 mg of frozen liver), total hepatic HYP was calculated (total liver, as obtained by multiplying liver weights with relative hepatic HYP).

### Liver histology and Immunohistochemistry

The left lobe was fixed in 5 ml 4% PFA at 4°C for 2 days for paraffin embedding. Formalin-fixed, paraffin-embedded (FFPE) liver sections were stained with hematoxylin and eosin (H&E) for assessment of liver structures and inflammation. For assessment of hepatic fibrosis, FFPE sections were stained with Sirius Red (Sigma, 365548-5G). FFPE liver sections were incubated with primary antibodies against α-smooth muscle actin (αSMA) (Abcam, ab5694, 1:100), rat anti-F4/80 (BioRad, MCA497R, 1:100) or rabbit anti-CYP2E1 (Sigma, HPA009128, 1:100) to assess activated HSC, resident macrophages and pericentral hepatocytes, respectively. The slides were scanned shortly after the staining procedure using the bright field microscope BX41. Digital pathological analysis was performed using ImageJ (https://imagej.nih.gov/ij/) on an equal number of pictures per mouse (10-15 images) under constant magnification (10X).

### Preparation of cryosections and lipid droplet staining

Part of the median lobe was embedded in tissue-Tek (VWR, 25608-930) and kept at –80°C till cryosectioning. Cryo sections (5μm thickness) were fixed in 4% PFA for 15 min, then briefly washed with running tap water and 60% isopropanol. Then cryosections were incubated with Bodipy (Life Technologies, D-3922, 1:250) for 30 min After rinsing steps with 60% isopropanol, cryosections were incubated for 5 min with Draq5 (Cell Signaling Technology, 4084L, 1:5000). Two tile scans of 9 images each per mouse for quantification were acquired using confocal microscopy (Leica SP8, UMM-Core facility Mannheim, Germany). Lipid droplets quantification was performed using ImageJ (https://imagej.nih.gov/ij/) on tilescans.

### Statistical analyses

Statistical analyses were performed in Prism (Version 8, GraphPad Software). Data are shown as mean ± SEM of 3-6 mice per group and two-tailed Student’s t test was calculated when shown. p-values <0.05 (*), < 0.01 (**), < 0.001 (***), < 0.0001 (****) are indicated.

## RESULTS

### Liver response to repeated toxic injuries switches at six weeks of exposure

To investigate the dynamics of liver responses to injuries induced by repeated application of a toxic chemical, we established a mouse model with repetitive CCl_4_ injections over ten weeks (Figure 1a) and analysed blood and liver samples harvested every week. For the control samples, we used both mice treated with oil for one and ten weeks. As shown in the survival analysis, in the first six weeks of CCl_4_ exposure per week 5% of mice died, which resulted in total loss of 10% of the animals by week six. Interestingly, after week six of exposure no further mortality caused by repeated CCl_4_ injuries was observed (Figure 1b). During the first six weeks, mice treated with CCl_4_ showed a significant gain in the liver to body weight ratio (Supporting figure 1a and b). Further, during this exposure period alterations in the survival rate and liver weight in these animals were accompanied by an increase of serum ALT (Figure 1c) (p < 0.05) and AST (Figure 1c) (p < 0.05), which are enzymes elevated in the blood in case of liver injury. No further increase of these values was detected after week six. Blood triglyceride (TG) levels continuously increased in a time-dependent manner until week five (Figure 1c) and returned to normal values towards week ten. No significant changes in blood cholesterol (Chol) was reported (Figure 1c). Markers for bile duct damage (alkaline phosphatase, ALP), and kidney function (Blood Urea Nitrogen, BUN) showed no major significant alterations (Supporting figure 1c-e) post week six. A trend for a decrease in glucose and total protein content was observed (Supporting figure 1f-g). These results indicate a switch of response program in the liver at six weeks of exposure to repetitive injuries from the phase of liver damage initiation and progression markers to a subsequent phase of adaptation leading to tolerance amongst the surviving mice.

### Time resolved tissue-based analysis divides the disease dynamics into initiation, progression and tolerance phases

To examine the dynamics of structural changes in the liver in response to repetitive injuries, we performed staining and quantification of samples from mice that were exposed to repetitive CCl_4_ administration for up to ten weeks. Compared to the control group, H&E staining of CCl_4_ treated livers demonstrated marked cell necrosis, inflammatory reaction and formation of septal fibrosis (Figure 1d, HE images). Moreover, we measured mRNA (Supporting figure 1h) and protein levels (Figure 1e) of CYP2E1, the key metabolizing enzyme of CCl_4_. Both mRNA and protein levels showed constant levels of CYP2E1 even after week six of repeated liver injury indicating stable metabolic capacity of the liver. Quantitative morphometric assessment of α-SMA positive (activated hepatic stellate cells; HSC) revealed massive accumulation during the first six weeks of repetitive CCl_4_ exposure (Figure 1f). No further increase in α-SMA positivity of HSC activation was observed at the later time points despite the continuous CCl_4_ administration (Figure 1f, α-Sma positivity). Likewise, the ECM deposition analyzed by the quantification of picrosirius red (PSR) positive areas showed the same behavior and did not increase beyond six weeks of exposure (Figure 1g). To ensure that these ECM-related changes are not local or sparse events, we used a biochemical assay to evaluate the hydroxyproline levels, HYP, a major component of collagen (Figure 1h), within the liver-tissue. With these analyses, we confirmed that the results obtained with quantitative morphometry did not only reflect local events but homogeneously distributed alterations caused by continuous CCl_4_ treatment. Our biochemical findings and histopathological features of the dynamics of fibrosis support the notion of an adaptive liver response that is able to tolerate repeated injuries after six weeks and is independent of the liver metabolic bioactivation of CCl_4_.

### Transcriptomics and proteomics divide the identified regulatory programs of liver fibrosis into three phases

The dynamics of structural changes detected at the tissue-level are indicative of underlying molecular alterations, which are best studied at the transcriptional and proteomic level. With these studies, we aimed at elucidating which regulatory program enables the liver to tolerate repeated injuries. As many molecular changes associated with fibrosis may overlay effects in the early adaptation phase, we profiled the transcriptome of the liver of 18 mice across six time points of our exposure series using Affymetrix microarrays. We validated the microarray derived transcriptomic data via RT-PCR of seven known fibrogenic genes, namely Col-1 α1, Col-1α2, α-Sma (Acta2), Timp1, Ctgf, Tgfβr1, and Tgfβ2 (Supporting figure 2a and b). A significant (p < 0.05) and positive correlation (mean ~0.72) between RT-PCR and microarray derived data was observed (Supporting figure 2a and b) for all tested genes. In the transcriptome, volcano plots showed the differentially regulated genes in 2, 6, and 10 weeks compared with control livers (Figure 2a). We also identified, per time point, the differentially expressed (DE) genes (FDR < 0.05) (Figure 2b). The largest differences were obtained at week six, comprising 1812 DE genes, which is in line with the tissue-level analysis. These highly variable properties were also observed via Principal Component Analysis (PCA) (Supporting figure 3a). To identify common differentially expressed genes, venn diagrams were generated based on deregulated (down and upregulated) targets at 1-2 weeks (phase I), 4-6 weeks (phase II) and 8-10 weeks (phase III). As shown, we identified 467 genes commonly regulated across all three phases (Figure 2c). Most of those genes were assigned to ECM and inflammation pathways. In addition, 210 genes were only differentially expressed in phase III (Figure 2c). Reduced molecular regulation in phase III after week six reflected previous findings (Figure 1) suggesting to refer to this phase as injury tolerance. Annotation of overrepresented pathways based on differentially expressed genes showed that besides several metabolic pathways the ECM pathway is induced during the initiation and progression phase (Supporting figure 3) and stabilized or reduced later. Analysis of time-resolved expression of selected genes (Figure 2d) showed a dynamic increase of fibrogenic targets e.g. Acta2, Col1a1, Col1a2 until week 6 and decreased during the tolerance phase. However, induction of Fasn (lipogenic target) was reported during the tolerance state (Figure 2d). Heatmap of most deregulated genes was shown in Figure 2e.

**Figure 2:**
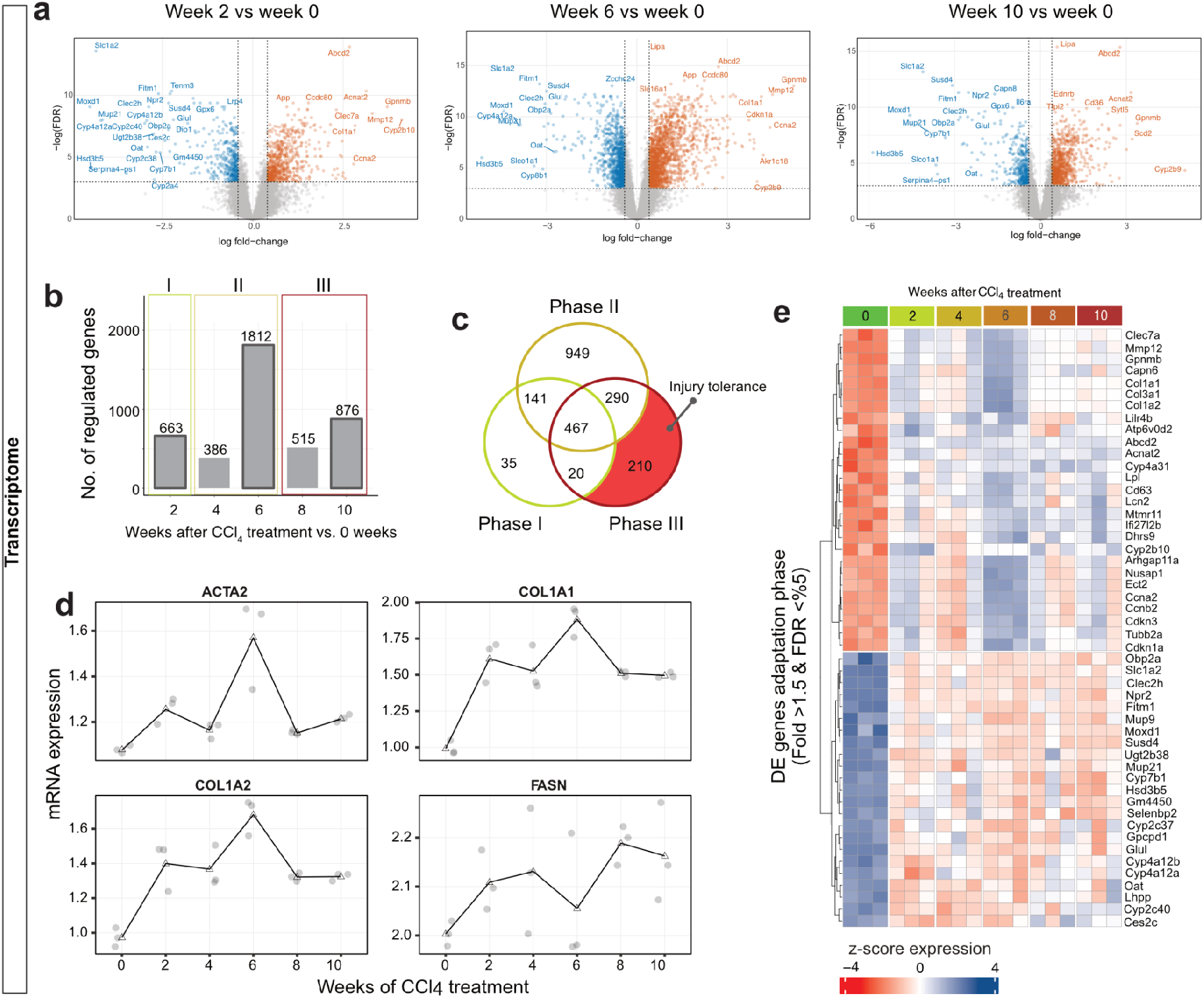
*Time-resolved transcriptome analysis of liver exposed to* CCl_4_. *(a) Volcano plots of DE genes in 2, 6 and 10 weeks compared with 0 week of* CCl_4_ *exposure. (b) The number of differentially regulated genes is visualized in the bar charts. Time points are grouped into three phases namely, initiation, progression and tolerance characterizing the disease course of liver fibrosis. (c) Venn diagrams show the unique and overlapped number of genes between the three phases. Interestingly, 210 genes were deregulated only in the tolerance phase. (d) show the regulation of selected genes in a time-resolved manner namely, ACTA2, COLaA1, COL1A2 and FASN. (e) Top deregulated genes during Phase III (tolerance) were visualized by heatmap. Transcriptomics data were obtained from 3 mice per time point*.

In addition, determinations of proteomic alterations were performed by multiplexing using isobaric labeling with TMT 10-plexes followed by MS/MS analysis, which resulted in the identification of in total 4222 proteins. For 2278 proteins, expression values were detected in all samples and further subjected to statistical analysis. Volcano plots showed the differentially regulated proteins in 2, 6, and 10 weeks compared with control livers (Figure 3a). Overall, similar dynamics as in the transcriptomics analysis was observed in proteomic data with 225 and 221 proteins differentially regulated in week four and six (Figure 3b). We performed PCA and differential expression analysis using linear models comparing each timepoint with the control time point to identify regulated proteins (Supporting figure 3b).

**Figure 3:**
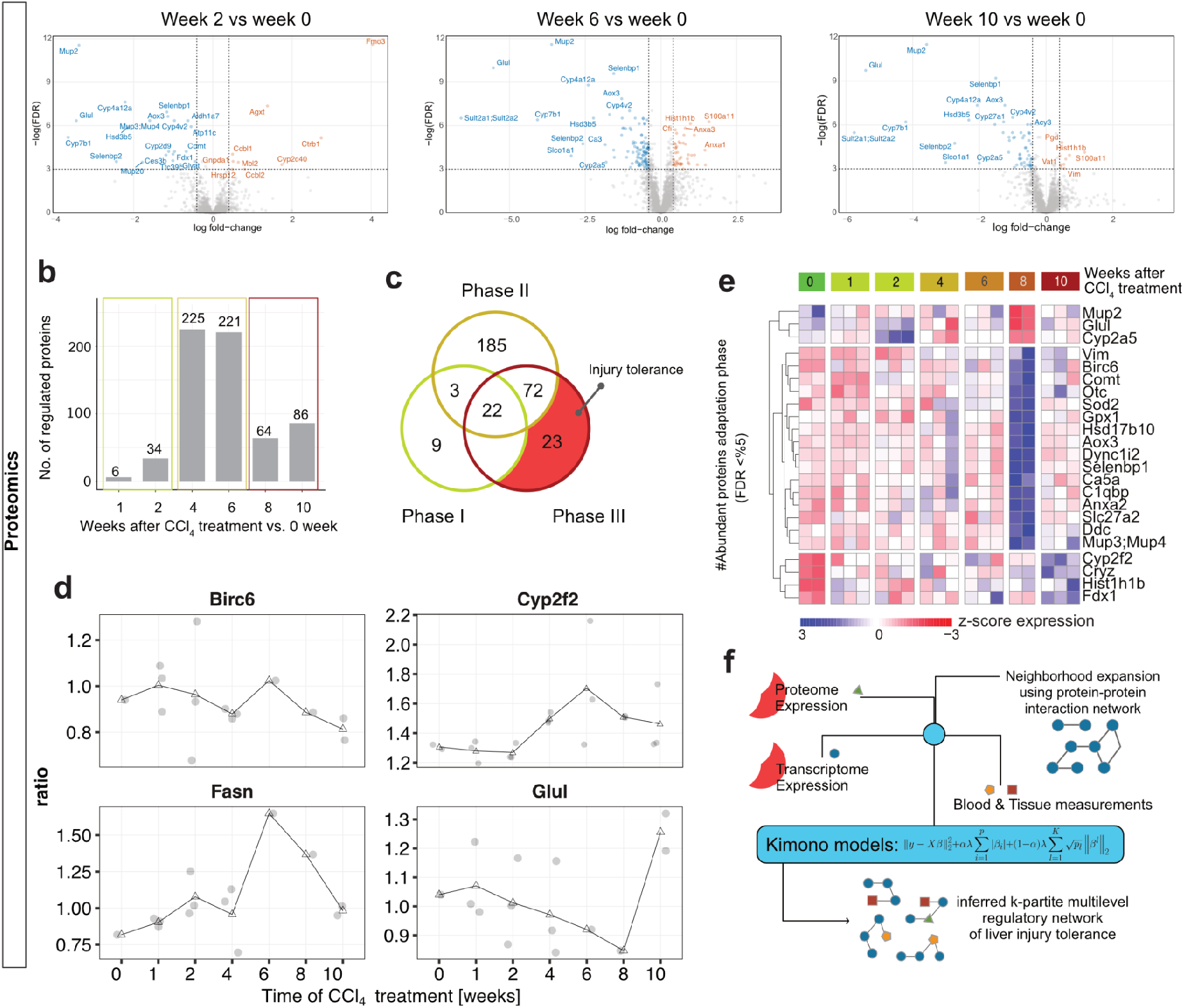
Time-resolved proteomic analysis of liver exposed to CCl_4_. *(a) Volcano plots of DE proteins in 2, 6 and 10 weeks compared with 0 week of* CCl_4_ *exposure. (b) The number of differentially regulated proteins is visualized in the bar charts. Time points are grouped into three phases namely, initiation, progression and tolerance characterizing the disease course of liver fibrosis. (c) Venn diagrams show the unique and overlapped number of proteins between the three phases. Interestingly, 23 proteins were deregulated only in the tolerance phase. (d) show the regulation of selected proteins in a time-resolved manner namely, Birc6, Cyp2f2, Fasn and Glul. (e) Top deregulated proteins during Phase III (tolerance) were visualized by heatmap. (f) Integration of proteomic, transcriptomic, blood and tissue measurements was performed by KiMONo models. Proteome data were obtained from 2-3 mice per time point*.

The number of regulated proteins compared to the amount of proteins identified by MS/MS was in line with the observed transcriptomic changes. To identify common differentially expressed proteins, Venn diagrams were generated based on deregulated (down and upregulated) targets at 1-2 weeks (phase II), 4-6 weeks (phase II) and 8-10 weeks (injury tolerance, phase III). As shown, we identified 22 common proteins (Figure 3c) across all analyzed time points. In addition, 23 proteins were only differentially expressed in the tolerance phase (Figure 3c). Furthermore, the heatmap showed 23 proteins were deregulated during the tolerance state (Figure 3e). Annotation of overrepresented pathways based on differentially expressed proteins showed that besides several metabolic pathways the ECM pathway is induced during the initiation and progression phase (Supporting figure 3). Analysis of time-resolved expression of selected proteins (Figure 3f) showed a dynamic increase of fibrogenic targets e.g. collagens, *Acta2* and stability of metabolic targets like *Cyp2f2* and *Glul* during the tolerance state. Taken together with the previous results obtained at the tissue level, the pattern of differentially expressed genes and proteins suggest that molecular regulation occurs in three phases: initiation (weeks 1-2) of immediate response patterns to repeated injuries, progression (weeks 4-6) of accumulating molecular changes upon repeated injuries and finally tolerance (weeks 8-10). The majority of initiation genes (95%) and proteins (75%) are also found in the progression phase. This led us to subsume the initiation and progression phase into the initial response program of liver injuries, which is in line with the survival, blood and tissue level analysis.

### Multi-level network of liver tolerance to repeated injuries captures human liver disease profiles

To dissect key factors determining the response of the liver to repeated injuries, we generated a fully integrated map combining transcriptomic, proteomic, biochemical and histopathological data. Briefly, we integrated and analyzed all available data using our recently published multi-omic network inference strategy – KiMONo (pipeline shown in Figure 3f, Figure 4). This allowed us to identify molecular patterns that covary with tissue-level histological parameters. In this inferred multi-level fibrosis network, nodes represent features like proteins, genes or biochemical parameters and connections denote statistically identified effects between them. This network was trimmed by excluding all node models with a low goodness of fit (R^2 < 0.1) as well as all effects weaker than beta < 0.002. The final network contained 8199 nodes connected via 16398 links (Supporting table 4). We identified tolerance phase specific network features via extracting nodes and modules associated with previously detected deregulated genes and proteins. This resulted in 68 injury-tolerant specific nodes and 13 distinct modules, with the smallest and biggest modules having 2 and 29 nodes respectively (Figure 5a and b). Small modules (2-3 nodes) were manually annotated to biological function using the GeneCards database (Stelzer et al. 2016), while the rest was annotated using the online pathway analysis tool PathwaX II (Ogris et. al. 2018). A total of 13 modules were identified during the tolerance state, of which 3 and 2 modules were related to lipid and carbohydrate metabolism (Figure 5a-b). The largest module was enriched for pathways in cancer (Supporting table 4 and 5, Module 11) encompassing key factors of TNFa, EGFR signaling and cell cycle. This observation triggered the comparison of the regulation of adaptation/tolerance phase (Figure 5a, b) with a variety of human diseases. Using eleven publicly available datasets on steatosis, NASH, ALD and HCC, we compared fold change of regulation of human homologs with the murine data (Supporting table 3, Figure 5c). Another large module was related to cancer pathways including 29 targets e.g. CDC42, CDC25, CDK5. Moreover, we detected several small modules, related to inflammatory and immune system pathways among the adaptation-specific modules. Here, consistent with human cohorts, GTF2IRD1, RXRA, JUN, N4BP2 and BCL3 were upregulated targets in inflammatory pathways. In conclusion, lipid and carbohydrate metabolic targets are predicted as modules that might play a role during the liver tolerance phase (Figure 4). We next focused on the known association of inflammation with disease progression (Supporting figure 5a and b). The results from analysis of F4/80 (marker for resident macrophages) stained livers and gene expression (*Adgre1*) revealed that F4/80 positive cells infiltrated in the injured liver (Supporting figure 5a and b) and further infiltration was observed during the tolerance phase. The detected levels of Adgre1 mRNA was in agreement with results obtained by IHC stained tissues (Figure 5a). These results demonstrated that the conspicuous pathways identified using KiMONo can be confirmed experimentally.

**Figure 4:**
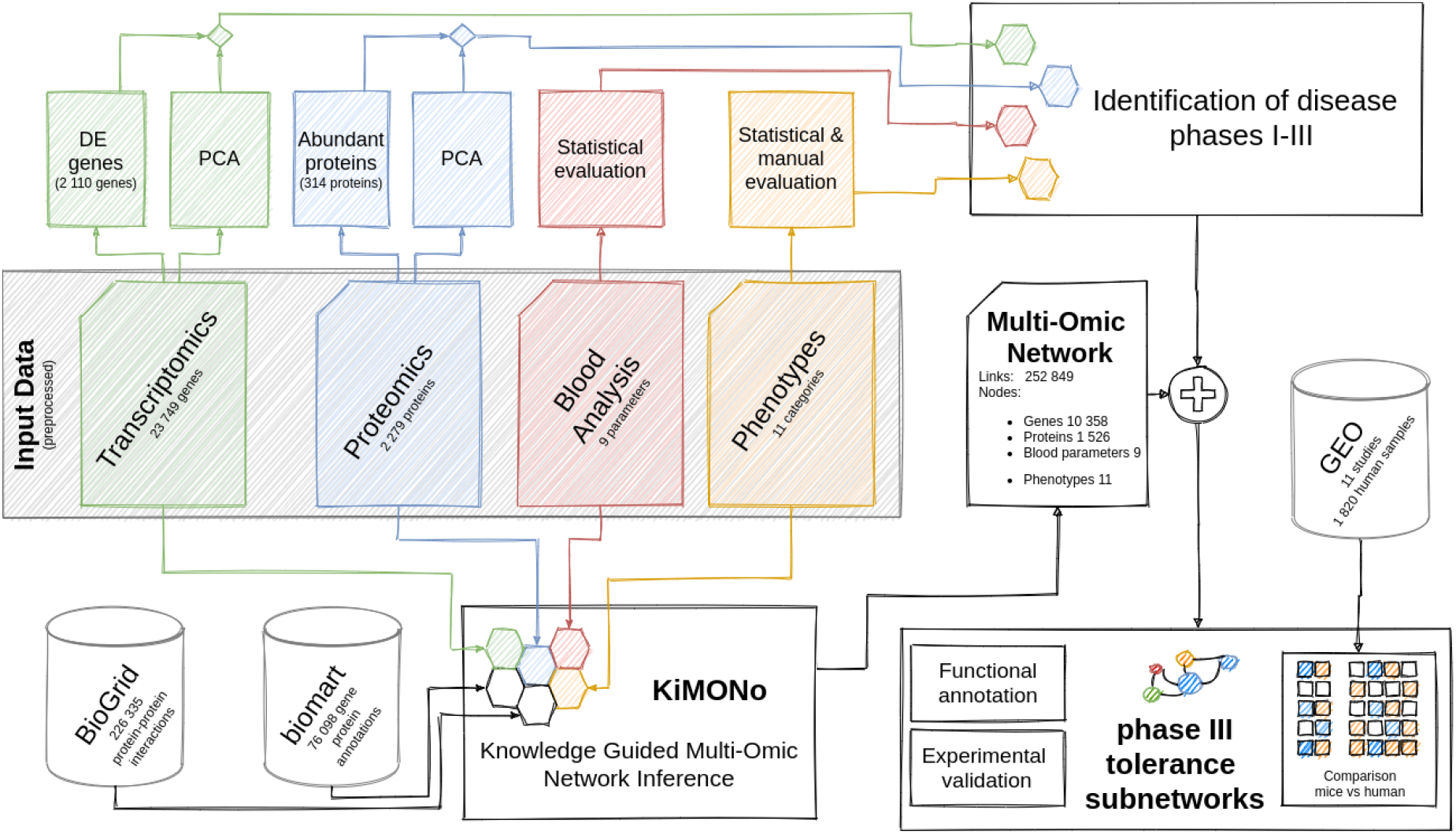
Data analysis workflow. The data sets generated via transcriptomics, proteomics as well as blood, histopathological and biochemical analysis were analyzed in an independent and integrative manner. The independent analysis was performed via state-of-the-art statistical approaches i.e. differential and consistently regulated genes and proteins. Combining the results, we identified three disease phases. Using the KiMONo algorithm,a fibrosis specific multi-omic network was inferred. Within this network we identified several nodes connecting tolerance phase specific features forming 13 distinct multi-omic modules. Some of these modules were experimentally validated and compared to 11 independent human studies of various liver diseases.

**Figure 5:**
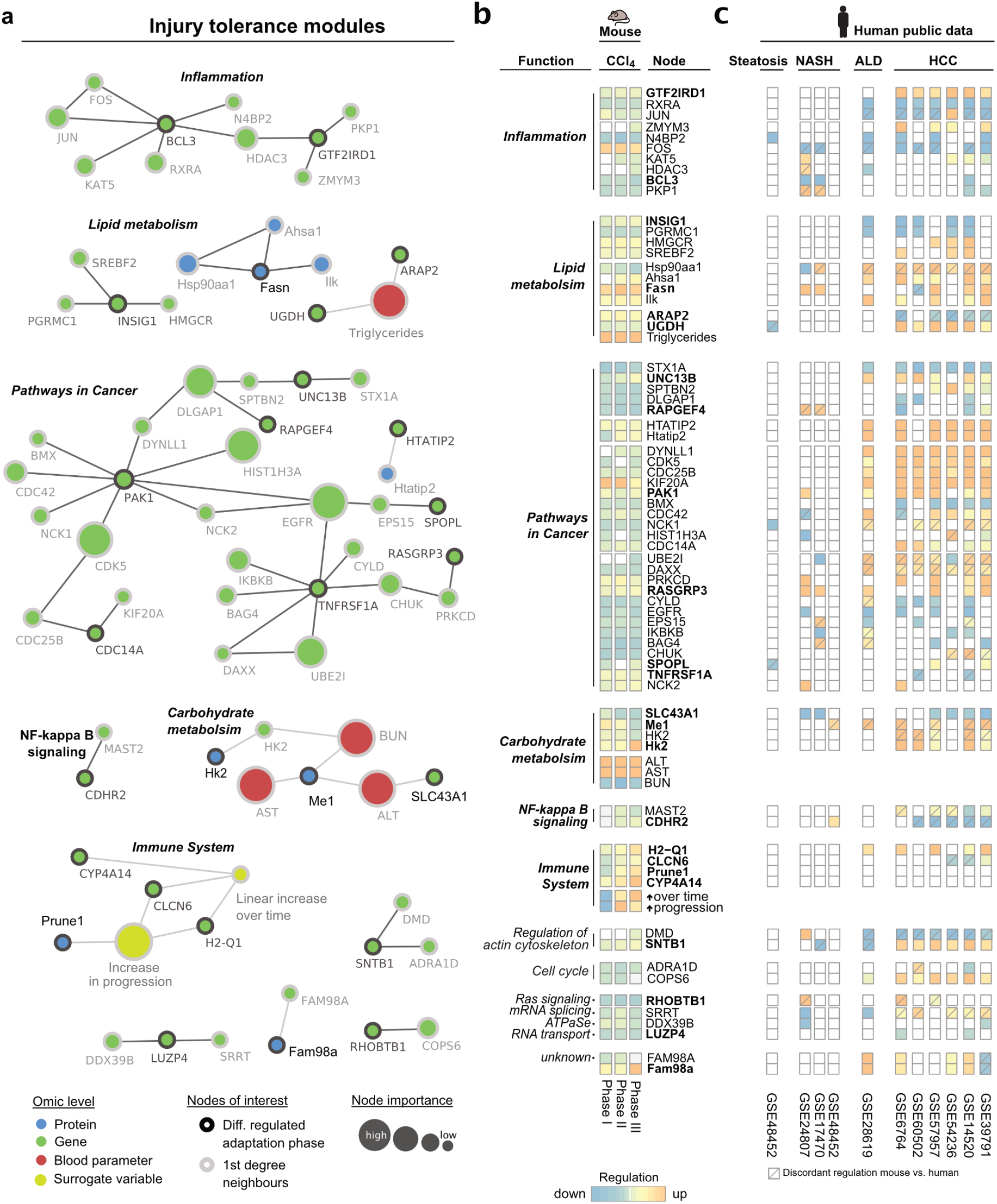
Tolerance specific modules in CCl_4_-induced fibrosis. *(a) Within the multi-omic fibrosis network we identified 13 tolerance phase specific modules by extracting differential regulated genes (green) and proteins (blue) and their 1st degree network neighbors. Network nodes are only connected when statistical effects are detected within the data. Node sizes refer to their importance within the network, which relate to high or low effects of* CCl_4_ *treatment. (b) Functional annotation and average regulation of network nodes for initiation, progression and tolerance phase. Significant (FDR < 0.05) downregulation (blue) and upregulation (red) are visualized within the heatmap. Bold node names denote uniquely differential regulation within the tolerance phase. (c) Significantly (FDR < 0.05) differential expressed genes of 11 human studies investigating steatosis, NASH, ALD and HCC*.

### Accumulation of lipid droplets resulting from increase of lipid metabolism in phase of toleration to repeated injuries

Lipid metabolism was specifically altered in the tolerance phase upon injuries induced by repeated CCl_4_ exposure. Therefore, we used IF, IHC and RT-PCR to confirm these findings. Firstly, we observed voids in CCl_4_ exposed liver tissue using HE, PSR, α-SMA (Supporting figure 5a). By histopathological analysis we observed that these voids were overloaded-hepatocytes with lipid droplets (Supporting figure 5b).

The presence of steatotic hepatocytes was in line with the KiMONo prediction. Therefore, we established Bodipy staining of lipid accumulation in the liver cells using cryosections and analyzed intracellular lipid accumulation using Bodipy positivity in liver cells. Interestingly, the results showed an increased intracellular lipid accumulation in a time-dependent manner (Figure 6a). Quantification of lipid droplets (Bodipy positive areas) revealed further lipid accumulation (Figure 6a) after six weeks in agreement with KiMONo prediction (Figure 5a). Furthermore, gene analysis of key regulators of lipid metabolism were quantified by RT-PCR. Fatty acid synthase (*FASN*), sterol regulatory element binding transcription factor 1 (*Srebp-1c*) and stearoyl-CoA desaturase 1 (*SCD-1*) as central regulators of de novo lipogenesis were significantly upregulated (Figure 6b) in eight and ten weeks of repeated CCl_4_ exposure (tolerance phase). The same results were observed in CCl_4_-induced fibrosis and Stellic animal model (steatosis-NASH based model as a positive control for lipid droplet recognition) (Figure 6c and d). The expression of carnitine palmitoyltransferase I (CPT1; a key enzyme of mitochondrial β-oxidation), acyl-CoA oxidase 1 (ACOX1; a key enzyme of peroxisomal β-oxidation) and peroxisome proliferator-activated receptors (PPAR-α and γ) was downregulated during the damage/regeneration phase and recovered to basal level during tolerance phase (Supporting figure 5c). This indicated that the lipid metabolism is disturbed upon repeated CCl_4_ exposure mainly during the tolerance phase. This de novo lipogenesis in the tolerance phase might lead to increased resistance of liver cells to repeated toxin exposure.

**Figure 6:**
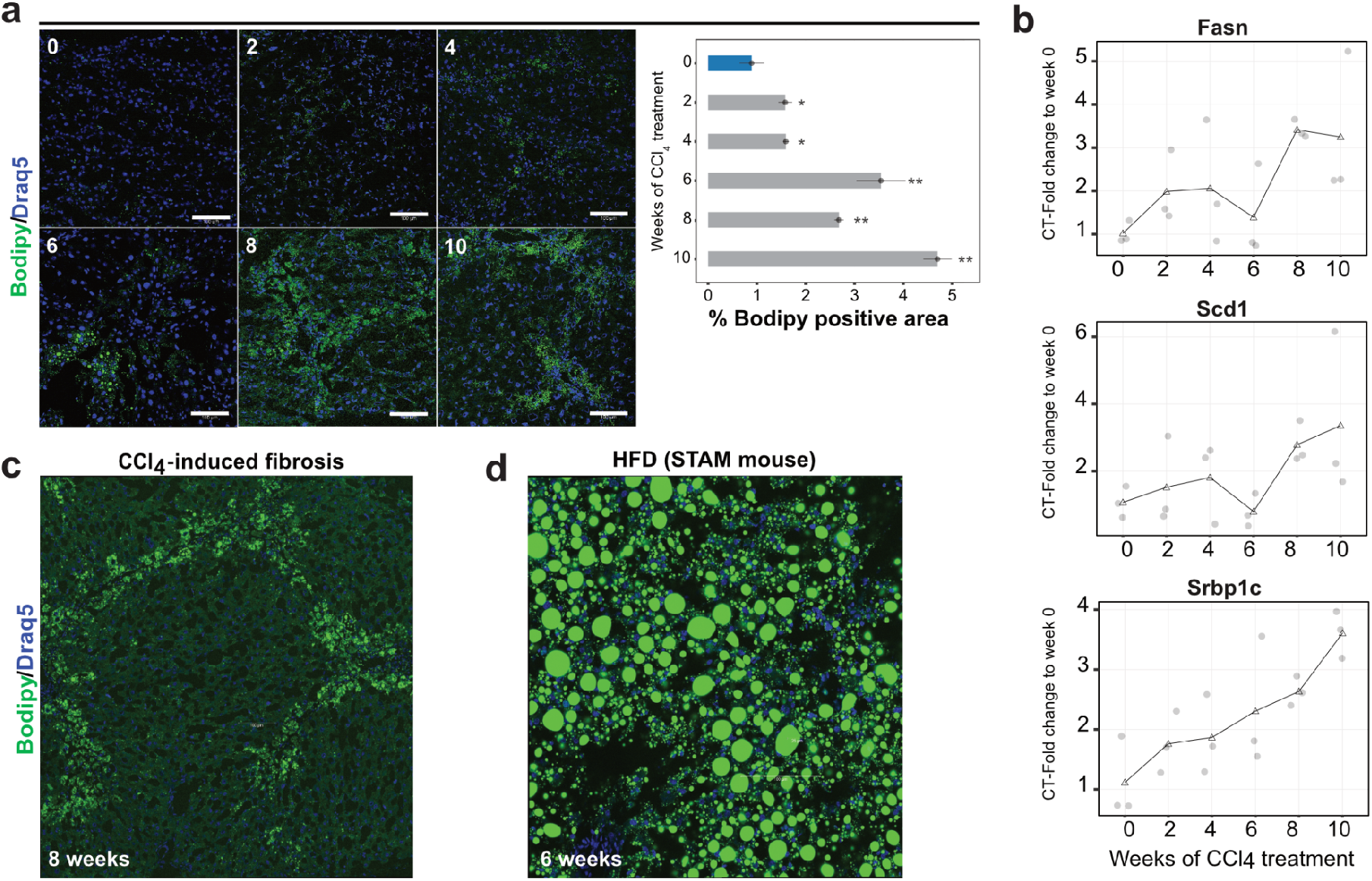
Lipid metabolism is induced during the adaptation phase as predicted by KiMONo integration analysis. *a) Bodipy staining to visualize and analyze lipid droplet accumulation in a time-resolved manner. Using a specific lipid droplet staining, namely Bodipy, we show that these voids are hepatocytes overloaded with lipid droplets. Scale bars are 100μm. b) mRNA levels of lipid metabolism-related targets i.e. Srbp1c, Scd1 and Fasn are analyzed by RT-PCR. Results were expressed as the mean of 3-6 mice ± SD, and were compared by two-way ANOVA test. *p <0.05 compared to 0 week (control). n=3-6 per group. c) and d) lower magnification images are shown from* CCl_4_-*induced fibrosis and Stellic animal model (steatosis-NASH based model as a positive control for lipid droplets recognition)*.

## DISCUSSION

Reversing liver fibrosis requires dissolving fibrous tissue and enhances quiescence of non-parenchymal cells as well as initiation of hepatocyte cell division. In this study, by a systematic biological analysis of high-throughput data of liver transcriptomes, proteomes, and clinical and histopathological parameters, we characterize week six as a transition time point critical for significant fibrosis development upon repeated CCl_4_ exposure. To avoid any interlobar difference, the same liver lobe from all tested mice was used to investigate downstream analysis. Therefore, the study did not consider lobe variations. Non-linear, however, significant accumulation of fibrotic tissue occurs and corresponds to METAVIR stage 2 or 3 during fibrogenesis in the first six weeks of CCl_4_. This result was in line with a previous report that an increase in fibrosis METAVIR stage is associated with a progressive increase in the fibrosis area, and the increase of fibrous tissue accumulation is not linear (Bedossa et al. 2003). Tissue-based analysis was correlated with other clinical parameters i.e. liver enzymes. Surprisingly, survival analysis revealed that the critical transition is week six. CCl_4_ is metabolized by CYP2E1 to form reactive trichloromethyl free radical and the trichloromethyl peroxyl radical (Weber et al. 2003). This critical transition in liver response is independent of CYP2E1 expression. Further, long-term downregulation of CYP2E1 and other CYPs are confirmed by Ghallab and co-workers (Ghallab et al. 2019) after CCl_4_ injections. Our results provide further insights for clinically and reliably signatures of mild and moderate fibrosis in order to achieve a complete fibrosis reversion. A limitation of this study is that we do not confirm on protein level identified targets in human fibrotic livers. Furthermore 10% of the mice were unable to cope with acute injury of CCl_4_ even though they are genetically identical. This might occur due to the interindividual responder effects, which are yet unexplored and unexplained.

To deepen our understanding on liver response, transcriptomics and proteomics are performed from the same mouse liver upon repeated CCl_4_ injections. This means tissue analysis is complemented with transcriptomics and proteomics data from the same mouse. Single layer analysis reflects a switch in number and differential expressed genes and proteins indicating different liver responses. However, the multi-level extensive profiling of a complex process, together with the employment of fully integrative approaches have the potential to unravel today’s overlooked features. Single omics have the tendency to result in several directions to follow up, and might have many bystanding effects. Pathway enrichment analysis annotations particularly those involved in fibrogenesis i.e. ECM and metabolism are in agreement with findings reported by Tuominen and co-workers (2021). To be considered, the dose of CCl_4_ was 0.8 g/kg and the mouse was sacrificed after 3 days of last injection in Tuominen et al. work (2021). Another transcriptomics study shows that no clear alterations of liver response i.e. ECM production and CYPs are between 2 and 6 months (tolerance phase) compared with 12 months after CCl_4_ administration (Ghallab et al. 2019). In this report, mice received 1 g/kg of CCl_4_ and were sacrificed 6 days after the last injection (Ghallab et al. 2019). In a rat study, Dong and colleagues report induction of ECM targets and metabolic pathways by proteomic and transcriptomics analysis of 9 weeks exposed rats for 1 ml/kg of CCl_4_ (Dong et al. 2016). However, the dynamic of the liver response is not studied by Dong and co-workers (2016) and also by several other reports (Nussler et al. 2014; Ghafoory et al. 2018; Gong et al. 2018). Overlaying evidence on multiple levels allows us to capture the major regulatory mechanisms. The overall CCl_4_ injury map has the potential to shed light on early injury effects but also mid-phase effects, where first-responder injury processes, like wound healing (ECM accumulation) are highly interesting effects to be further investigated. This multilevel analysis indicates a point where the injured organ, the liver, cannot cope with excessive ECM accumulation and switches the program to be more tolerant to further damage. This is a well-known toxicological response called drug tolerance.

KiMONO integration predicted 13 modules to be deregulated in the tolerance phase; among those cancer pathways, lipid metabolism and inflammation targets were included. Series of validation experiments for lipid metabolism reveal accumulation of microvesicular droplets in livers of eight and ten weeks. At these time points, several lipogenic targets i.e. Srebp1c, Scd1 and Fasn, are significantly induced compared to control and six weeks mice. It has been shown that PPARα is a key protein involved in liver lipid metabolism (Kersten 2014) and its induction results in de novo hepatic lipogenesis (Oosterveer et al. 2009). However, in our study both PPARα and γ are not significantly induced in tolerance compared with initiation and progression phase. Moreover, as predicted, sustained inflammatory processes by maintaining F4/80 cells in the vicinity of fibrosis are validated. Therefore, in CCl_4_ -exposed mice after six weeks it seems that reversion of TG in the blood to normal level and accumulation of microvesicular droplets are features of this liver tolerance. Careful analysis of the predicted modules in the tolerance phase identified several targets that were consistently deregulated in human liver diseases. Although our study has identified high confidence candidate targets for fibrosis tolerance, follow-up with knockdown/overexpression in specific cell types in the liver will be required for functional validation. In conclusion, in the present study we report a list of tolerance-related modules generated by unbiased multi-level analysis of time-resolved CCl_4_-exposed mice. Despite CCl_4_ being well-established hepatotoxic for regeneration and fibrosis, fatty liver is not carefully reported and studied. KiMONO prediction followed by experimental validation indicates that at a transition point the liver is not able to cope with repeated damage and initiates a tolerance program by sustained inflammation and accumulation of small lipid droplets. Further experiments are required to mechanistically investigate these protective modules for therapeutic activation.

## ACKNOWLEDGEMENT

The authors acknowledge the support of the Core Facility for Imaging and microarray unit Mannheim, Medical Faculty Mannheim, Heidelberg University; Marvin Wäsch for the technical support for the proteome sample preparation and measurements.

## AUTHOR CONTRIBUTION

SH, AO, PE, YG and WP conceived animal studies, blood analysis, RNA isolation, RT-PCR, IHC and IF staining and image analysis. IS, BH, LD performed proteomics measurements supervised by UK. SH, CO and WS performed transcriptomics analysis. NM performed the proteomics analysis. CO performed multi-omic data integration and multi-modal network analysis, comparison to human data, pathway analysis as well as data visualization. SH, CO and NM wrote the manuscript. UK, and SD performed critical revision of the manuscript. UK, SH, SD, JH, FJT, CO and NM acquired funding for this study. All authors read the final version of the manuscript. SH and NM designed the experimental and bioinformatics analysis strategy, supervised the study and interpreted the results.

