## supporting for "Repeated toxic injuries of murine liver are tolerated through microsteatosis and mild inflammation"

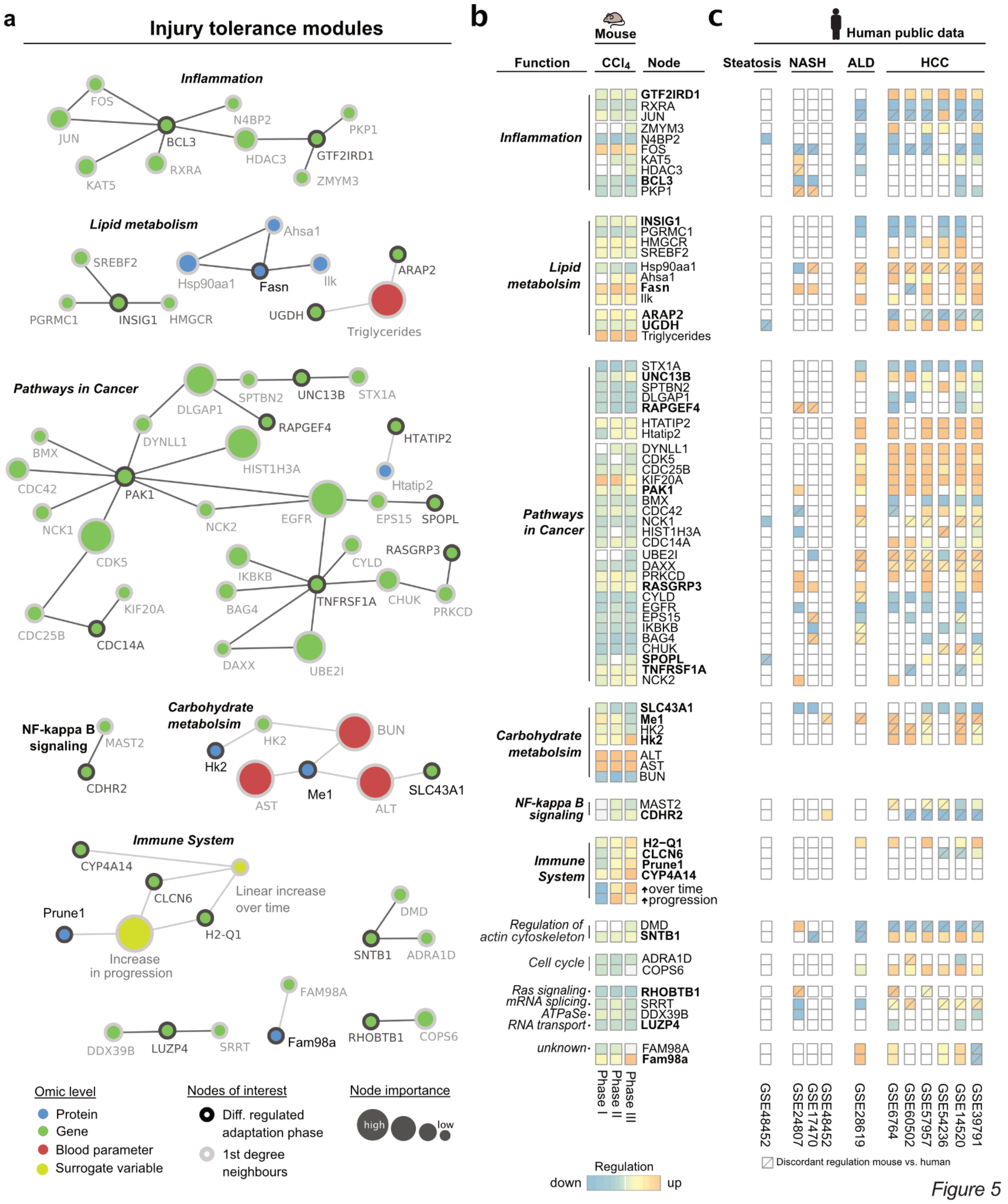


***Figure 5: Tolerance specific modules in $CCl_4$-induced fibrosis.*** *(a) Within the multi-omic fibrosis network we identified 13 tolerance phase specific modules by extracting differential regulated genes (green) and proteins (blue) and their 1st degree network neighbors. Network nodes are only connected when statistical effects are detected within the data. Node sizes refer to their importance within the network, which relate to high or low effects of* $CCl_4$ *treatment. (b) Functional annotation and average regulation of network nodes for initiation, progression and tolerance phase. Significant (FDR < 0.05) downregulation (blue) and upregulation (red) are visualized within the heatmap. Bold node names denote uniquely differential regulation within the tolerance phase. (c) Significantly (FDR < 0.05) differential expressed genes of 11 human studies investigating steatosis, NASH, ALD and HCC.*

**Accumulation of lipid droplets resulting from increase of lipid metabolism in phase of toleration to repeated injuries**

Lipid metabolism was specifically altered in the tolerance phase upon injuries induced by repeated $CCl_4$ exposure. Therefore, we used IF, IHC and RT-PCR to confirm these findings. Firstly, we observed voids in $CCl_4$ exposed liver tissue using HE, PSR, α-SMA (Supporting figure 5a). By histopathological analysis we observed that these voids were overloaded-hepatocytes with lipid droplets (Supporting figure 5b).

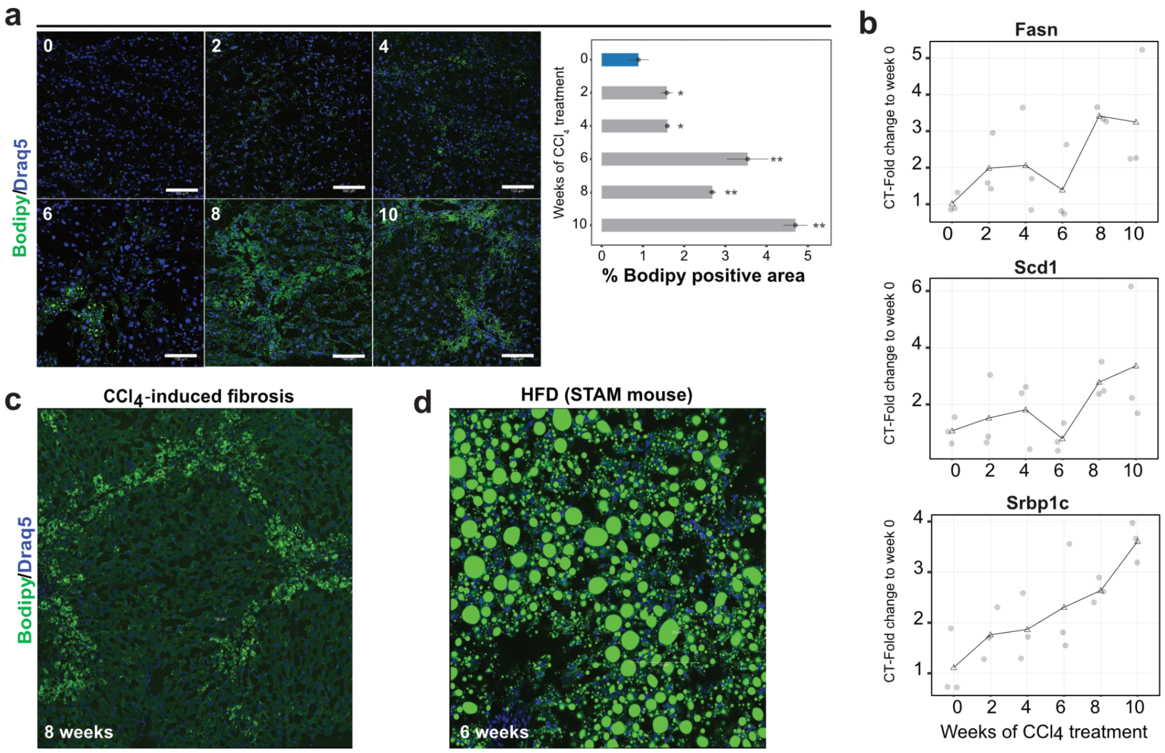


***Figure 6: Lipid metabolism is induced during the adaptation phase as predicted by KiMONo integration analysis.*** *a) Bodipy staining to visualize and analyze lipid droplet accumulation in a time-resolved manner. Using a specific lipid droplet staining, namely Bodipy, we show that these voids are hepatocytes overloaded with lipid droplets. Scale bars are 100µm. b) mRNA levels of lipid metabolism-related targets i.e. Srbp1c, Scd1 and Fasn are analyzed by RT-PCR. Results were expressed as the mean of 3-6 mice ± SD, and were compared by two-way ANOVA test. *p <0.05 compared to 0 week (control). n=3-6 per group. c) and d) lower magnification images are shown from* $CCl_4$*-induced fibrosis and Stellic animal model (steatosis-NASH based model as a positive control for lipid droplets recognition).*

#### FUNDING

This study was supported by the BMBF (German Federal Ministry of Education and Research) Project LiSyM (Grant PTJ-FKZ: 031L0043; 031L0042; 031L0047) and funding for NSM. Project TRY-IBD Grant 01ZX1915B for CO.

#### AUTHOR CONTRIBUTION

SH, AO, PE, YG and WP conceived animal studies, blood analysis, RNA isolation, RT-PCR, IHC and IF staining and image analysis. IS, BH, LD performed proteomics measurements supervised by UK. SH, CO and WS performed transcriptomics analysis. NM performed the proteomics analysis. CO performed multi-omic data integration and multi-modal network analysis, comparison to human data, pathway analysis as well as data visualization. SH, CO and NM wrote the manuscript. UK, and SD performed critical revision of the manuscript. UK, SH, SD, JH, FJT, CO and NM acquired funding for this study. All authors read the final version of the manuscript. SH and NM designed the experimental and bioinformatics analysis strategy, supervised the study and interpreted the results.

### Supporting Figures

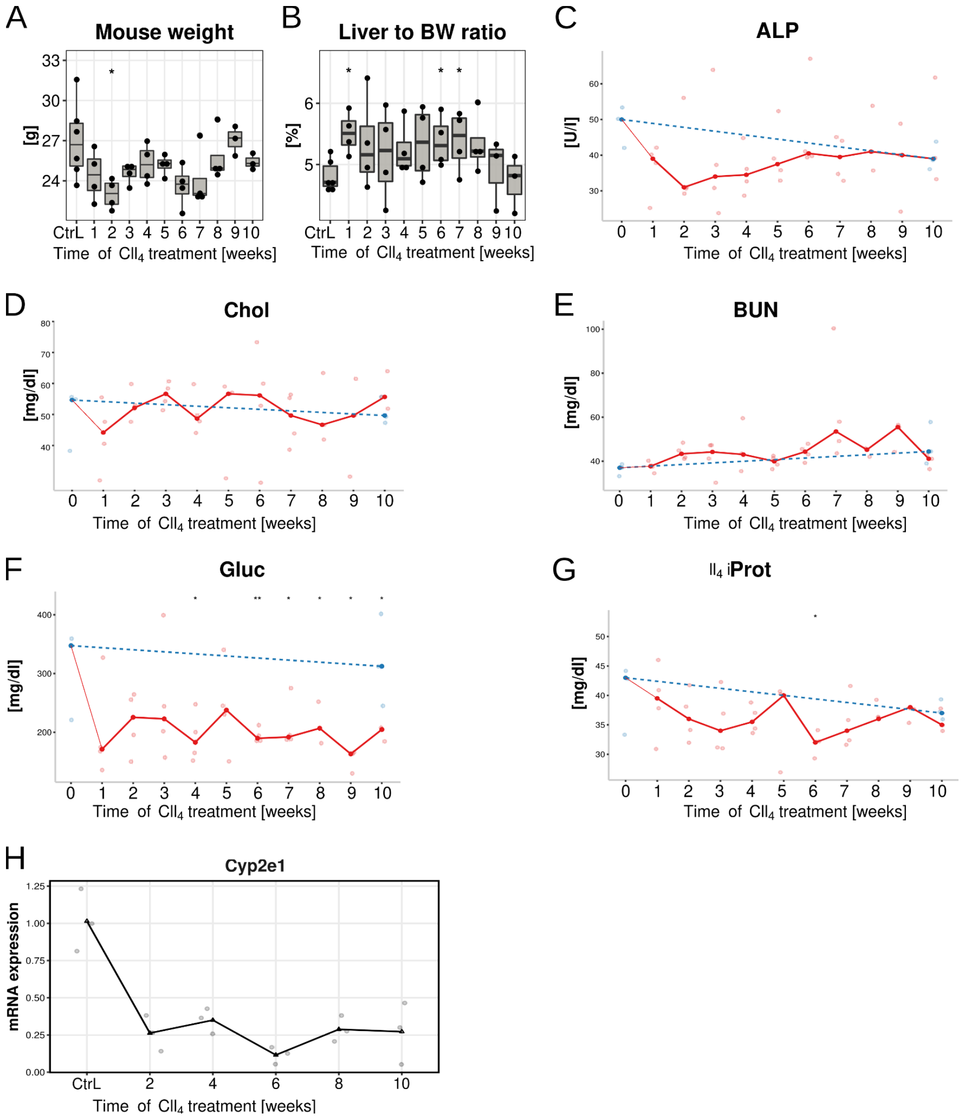

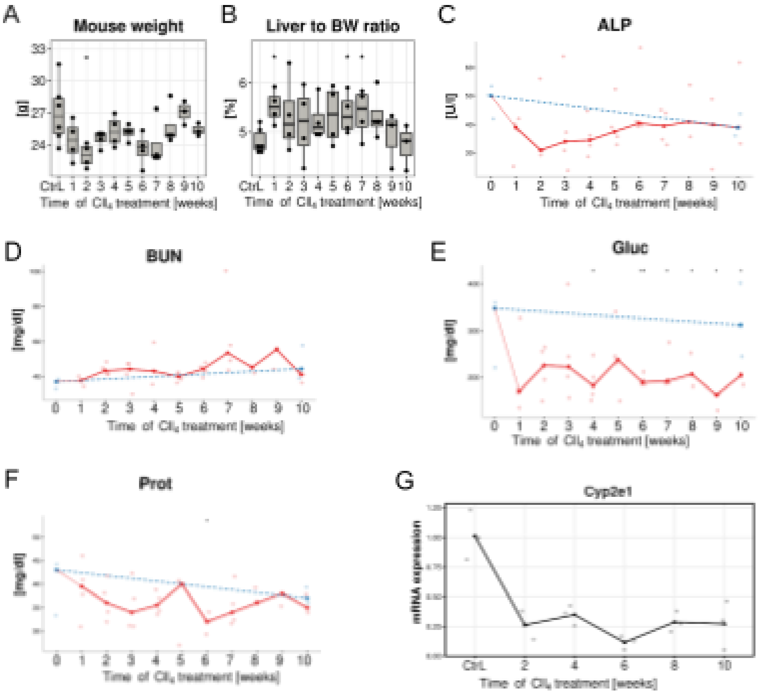


***Supporting Figure 1: Mouse and liver weight and metabolic parameters during fibrogenesis.*** *(A) Mouse weight. (B) Liver weight as a ratio of body weight. (C-F) Time resolved measurements of ALP, BUN Gluc, and total protein levels in blood of mice exposed to $CCl_4$. (G) mRNA level of Cyp2e1 analyzed by RT-PCR. Results are shown as mean ± SD, and were compared by two-way ANOVA test. *p <0.05 compared to 0 week (control). n=3-6 per group. $CCl_4$, carbon tetrachloride; ALP, alkaline phosphatase; BUN, blood urea nitrogen; Gluc, glucose; Prot, total protein; Cyp2e1, Cytochrome P4502e1.*

A

| Gene name | Gene symbol | r | p |
|---|---|---|---|
| Collagen 1α1 | Col1α1 | 0.5056 | 0.0323 |
| Collagen 1α2 | Col1α2 | 0.8680 | <0.0001 |
| Alpha-smooth muscle actin | Acta2 | 0.9650 | <0.0001 |
| TIMP metallopeptidase inhibitor 1 | Timp1 | 0.9785 | <0.0001 |
| Connective tissue growth factor | Ctgf | 0.6971 | 0.0013 |
| Transforming growth factor beta receptor I | Tgfβr1 | 0.5253 | 0.0252 |
| Transforming growth factor-beta 2 | Tgfβ2 | 0.4854 | 0.0412 |

B

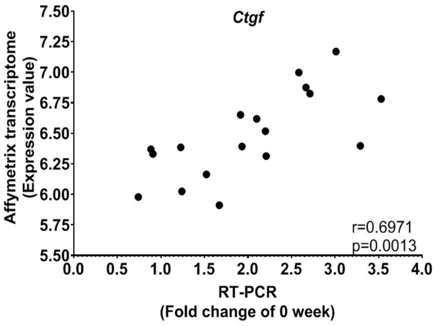


***Supporting Figure 2:*** *Analysis of 7 selected reference genes comparing RT-PCR and Affymetrix-based transcriptome in 18 mice. (A) Table shows Coefficient of Pearson correlation (r) and p values between each target analysed by RT-PCR and transcriptomics. (B) Connective tissue growth factor (Ctgf) result is blotted as an example of correlation between fold change of RT-PCR of 0 week and expression value based on transcriptome.*

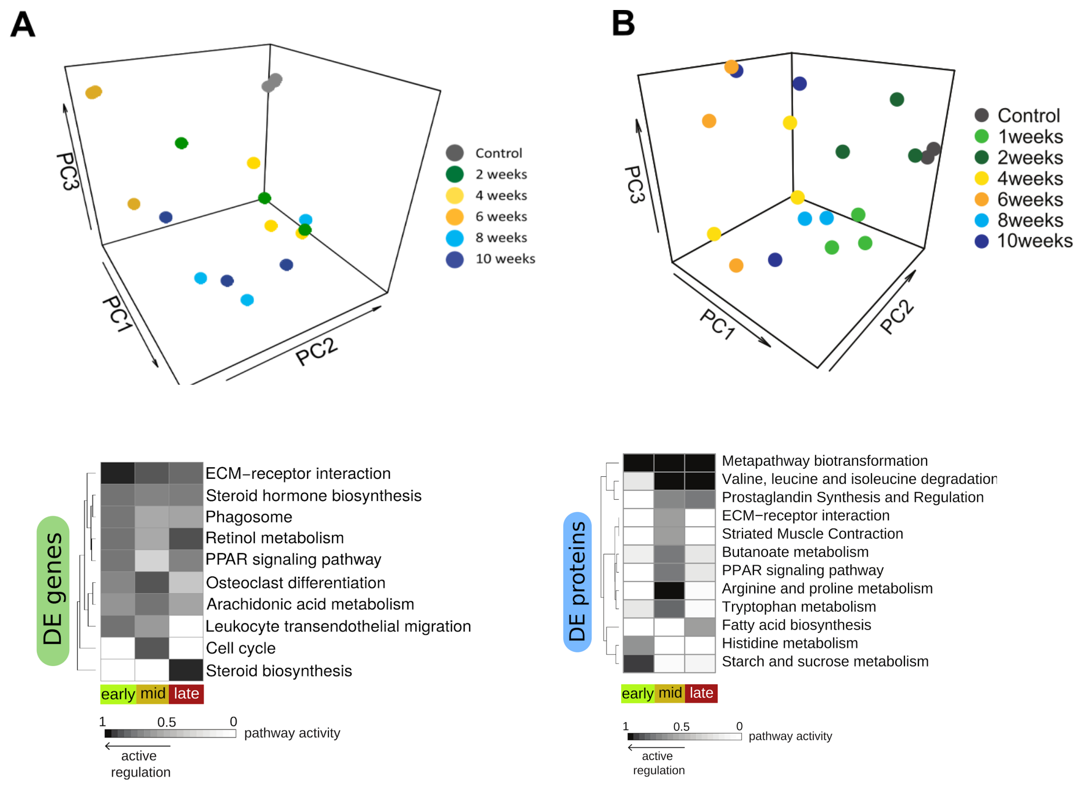


***Supporting Figure 3:*** *(A) PCA of the transcriptome across all time points. (B) PCA of the proteome across all time points. Pathway analysis of DE genes and abundant proteins across all three identified phases upon chronic $CCl_4$ injections.*

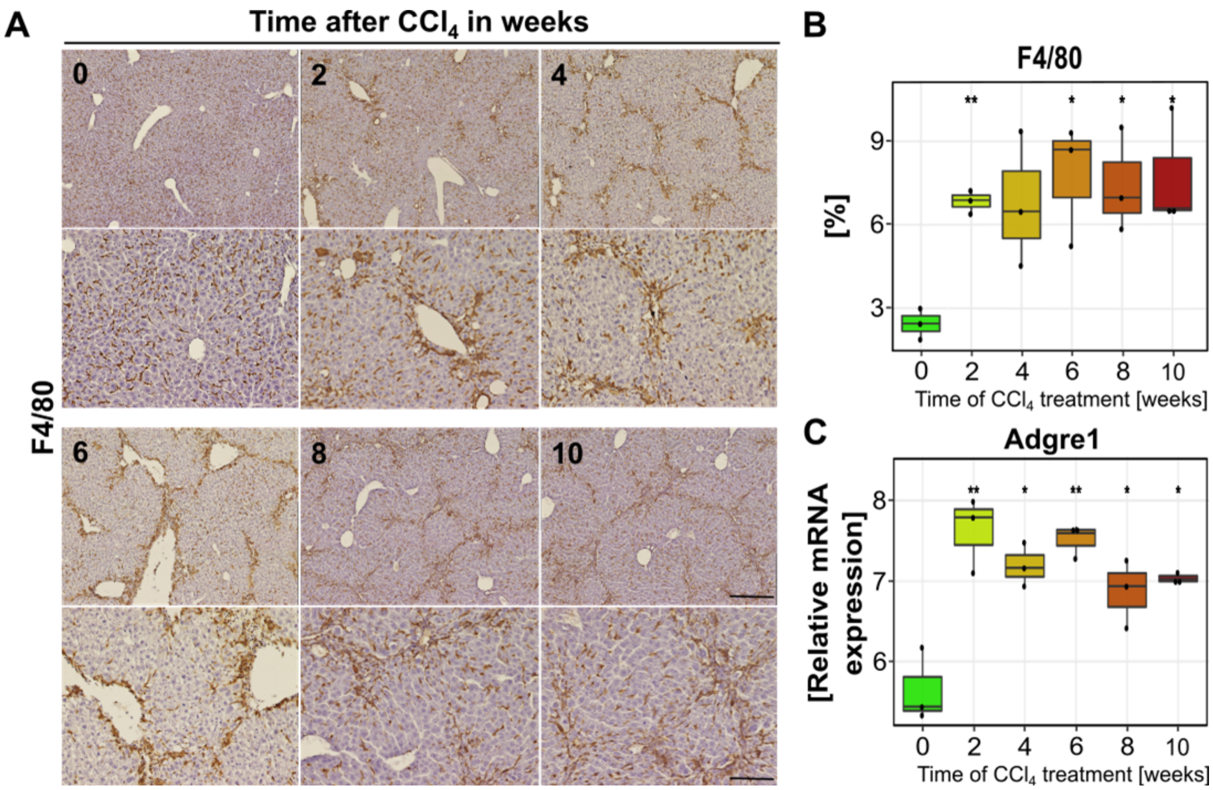


***Supporting Figure 4. F4/80 expression is sustained during the adaptation phase after repeated $CCl_4$ injections.*** *(A) Liver sections were prepared from liver tissue samples of control and mice with $CCl_4$ treatment for 10 weeks and subjected to immunostaining against F4/80. Scale bars are 100µm and 200µm for closeups and overview images, respectively. (B) Quantification of positive F4/80 signals from stained slides over the total area. (C) mRNA expression of F4/80 assigned gene (Adgre1) extracted from transcriptomic datasets. Results were expressed as mean ± SD, and were compared by two-way ANOVA test. *p <0.05, **p <0.01 compared to 0 week (control). n=3-6 per group.*

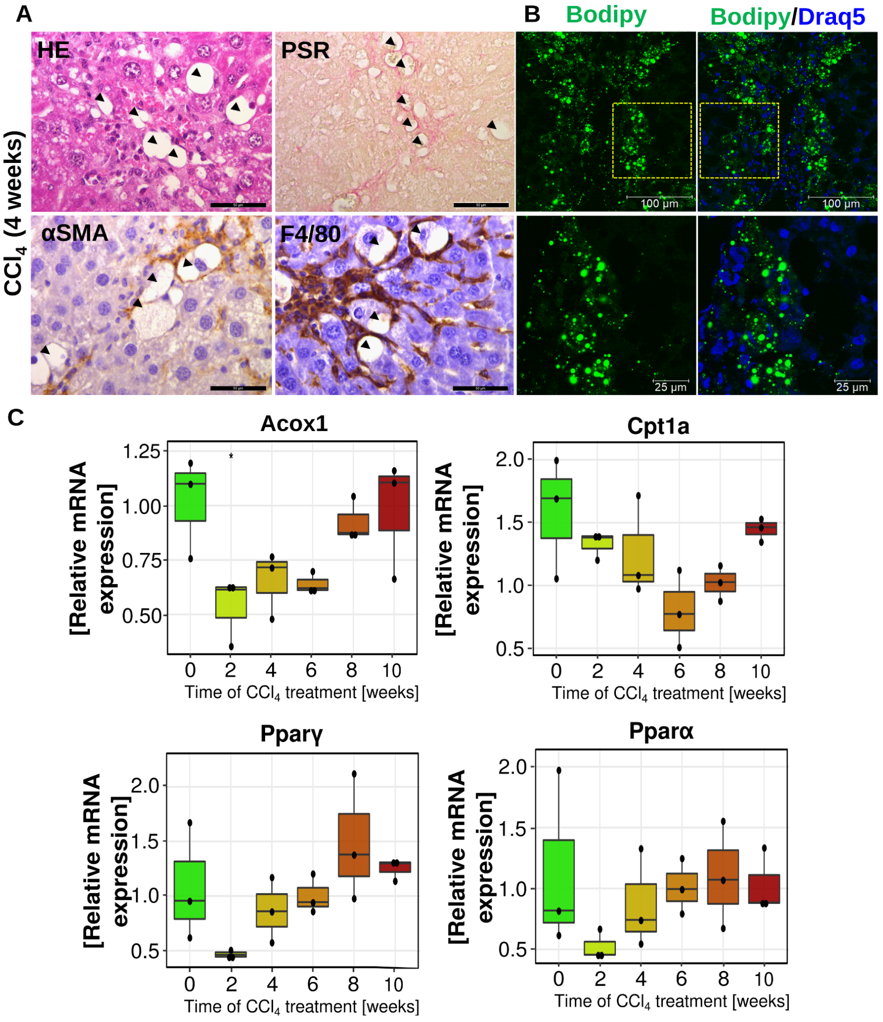


***Supporting Figure 5: Whitish voids in $CCl_4$-induced liver fibrosis are lipid droplets containing hepatocytes.*** *(A) Voids appeared as whitish areas in the liver tissue. The spatial distribution of these voids was along fibrotic regions. By HE and PSR, these voids have a specific circular structure and in some cases small nuclei are observed. Additionally, we found that these voids are surrounded by alpha-SMA or F4/80 positive cells similar to hepatic crown-like structures described for NASH. (B) Using a specific lipid droplet staining namely bodipy, we show that these voids are hepatocytes overloaded with lipid droplets. (C) mRNA levels of lipid metabolism-related targets i.e. Acox1, Cpt1a, Pparγ and Pparα are analysed by RT-PCR. Results were expressed as mean ± SD, and were compared by two-way ANOVA test. *p <0.05 compared to 0 week (control). n=3-6 per group.*

#### Supporting Tables

***Supporting Table 1:*** *RNA concentration and integrity isolated from oil and* $CCl_4$*-exposed mouse livers. This RNA was used for transcriptomics and RT-PCR analysis. 50ul was prepared (100ng/ul) by dilution in nuclease-free water Affymetrix (J71786 Water, Thermo Scientific). For Affymetrix transcriptomics 100 ng/ul (20ul) was used.*

| Mouse ID | Strain | Treatment | Time point | RIN | RNA conc (ng/ul) |
|---|---|---|---|---|---|
| Mouse 1 | C57/6-B6N | Oil | 1 weeks | 8.70 | 1489.4 |
| Mouse 2 | C57/6-B6N | Oil | 1 weeks | 8.30 | 2164.7 |
| Mouse 3 | C57/6-B6N | Oil | 1 weeks | 8.40 | 2481.4 |
| Mouse 1 | C57/6-B6N | $CCl_4$ | 2 weeks | 9.50 | 1816.8 |
| Mouse 2 | C57/6-B6N | $CCl_4$ | 2 weeks | 9.50 | 930.90 |
| Mouse 3 | C57/6-B6N | $CCl_4$ | 2 weeks | 8.90 | 1979.5 |
| Mouse 1 | C57/6-B6N | $CCl_4$ | 4 weeks | 8.20 | 3173.0 |
| Mouse 2 | C57/6-B6N | $CCl_4$ | 4 weeks | 8.80 | 1943.2 |
| Mouse 3 | C57/6-B6N | $CCl_4$ | 4 weeks | 7.80 | 2695.6 |
| Mouse 1 | C57/6-B6N | $CCl_4$ | 6 weeks | 8.20 | 2018.0 |
| Mouse 2 | C57/6-B6N | $CCl_4$ | 6 weeks | 8.50 | 2102.0 |
| Mouse 3 | C57/6-B6N | $CCl_4$ | 6 weeks | 9.10 | 1464.0 |
| Mouse 1 | C57/6-B6N | $CCl_4$ | 8 weeks | 8.90 | 3631.8 |
| Mouse 2 | C57/6-B6N | $CCl_4$ | 8 weeks | 9.00 | 2054.5 |
| Mouse 3 | C57/6-B6N | $CCl_4$ | 8 weeks | 9.00 | 731.80 |
| Mouse 1 | C57/6-B6N | $CCl_4$ | 10 weeks | 9.00 | 2532.1 |
| Mouse 2 | C57/6-B6N | $CCl_4$ | 10 weeks | 8.60 | 4241.1 |
| Mouse 3 | C57/6-B6N | $CCl_4$ | 10 weeks | 9.30 | 1863.9 |

***Supporting Table 2***. *Sequences of primer pairs used for real time PCR.*

| Gene | Forward | Reverse |
|---|---|---|
| *Acox1* | GAATTTGGCATCGCAGACCC | CGGGTGCATCCATTTCTCCT |
| *Acta2* | TTCGCTGTCTACCTTCCAGC | GAGGCGCTGATCCACAAAAC |
| *Col1α1* | GGAGAGAGCATGACCGATGG | AAGTTCCGGTGTGACTCGTG |
| *Col1α2* | AGTCGATGGCTGCTCCAAAA | AGCACCACCAATGTCCAG AG |
| *Cpt1a* | CGGACTCCGCTCGCTCATT | GGGAGGGGTCCACTTTGGTA |
| *Ctgf* | AGATTGGAGTGTGCACTGCCAAAG | TCCAGGCAAGTGCATTGGTATTTG |
| *Cyp2e1* | CGTTGCCTTGCTTGTCTGGA | AAGAAAGGAATTGGGAAAGGTCC |
| *Fasn* | ACAATGGACCCCCAGCTTCG | CAGACGCCAGTGTTCGTTCC |
| *Pparα* | TGCAGCCTCAGCCAAGTTGAA | GTTCCCGAACTTGACCAGCC |
| *Pparγ* | ACGTTCTGACAGGACTGTGT | CTGTGTCAACCATGGTAATTTCA |
| *Ppia* | GAGCTGTTTGCAGACAAAGTC | CCCTGGCACATGAATCCTGG |
| *Scd1* | AACAGTGCCGCGCATCTCTA | GAAGCCCAAAGCTCAGCTACTC |
| *Srbp1c* | GGAGCCATGGATTGCACATT | GGCCCGGGAAGTCACTGT |
| *Tgfβ2* | GCAGATCCTGAGCAAGCTG | GTAGGGTCTGTAGAAAGTGG |
| *Tgf-βr1* | GAACTGTTTTGATTGGCATC | AAGAAGGGACCTACACTATTT |
| *Timp1* | GGCATCTGGCATCCTCTTGT | ACTCTTCACTGCGGTTCTGG |

***Supporting Table 3.*** *Publically available Human GEO datasets.*

| GEO Accession # | Published | *Control | HCC | NASH | ALD | NAFLD | Obese | Steatosis |
|---|---|---|---|---|---|---|---|---|
| GSE14520 | Roessler et al. 2010 | 220 | 225 | - | - | - | - | - |
| GSE24807 | Liu et al. 2011 | 5 | - | 12 | - | - | - | - |
| GSE39791 | Kim et al. 2014 | 72 | 72 | - | - | - | - | - |
| GSE48452 | Ahrens et al. 2013 | 14 | - | 18 | - | 27 | 27 | 14 |
| GSE57957 | Mah et al. 2014 | 39 | 39 | - | - | - | - | - |
| GSE60502 | Wang et al. 2014 | 18 | 18 | - | - | - | - | - |
| GSE6764 | Wurmbach et al. 2007 | 10 | 10 | - | - | - | - | - |
| GSE17470 | Baker et al. 2010 | 4 | - | 7 | - | - | - | - |
| GSE28619 | Affò et al. 2013 | 7 | - | - | 15 | - | - | - |
| GSE54236 | Villa et al. 2016 | 80 | 81 | - | - | - | - | - |

***Supporting Table 4:*** *Network nodes representing mouse genes, and proteins, their subnetwork and the corresponding human ortholog.*

| Mus Musculus Node | Subnetwork | Homo Sapiens Ortholog |
|---|---|---|
| COPS6 | 1 | COPS6 |
| RHOBTB1 | 1 | RHOBTB1 |
| HTATIP2 | 2 | HTATIP2 |
| Q9Z2G9 | 2 | HTATIP2 |
| CDHR2 | 3 | CDHR2 |
| MAST2 | 3 | MAST2 |
| ARAP2 | 4 | ARAP2 |
| SNHG14 | 4 | |
| UGDH | 4 | UGDH |
| CLCN6 | 5 | CLCN6 |
| CYP4A14 | 5 | |
| H2-Q1 | 5 | HLA-A,HLA-B,HLA-C,HLA-D,HLA-E,HLA-F |
| Q8BIW1 | 5 | PRUNE1 |
| ADRA1D | 6 | ADRA1D |
| DMD | 6 | DMD |
| SNTB1 | 6 | SNTB1 |
| BCL3 | 7 | BCL3 |
| FOS | 7 | FOS |
| GTF2IRD1 | 7 | GTF2IRD1 |
| HDAC3 | 7 | HDAC3 |
| JUN | 7 | JUN |
| KAT5 | 7 | KAT5 |
| N4BP2 | 7 | N4BP2 |
| PKP1 | 7 | PKP1 |
| RXRA | 7 | RXRA |
| ZMYM3 | 7 | ZMYM3 |
| DX39B | 8 | |
| LUZP4 | 8 | LUZP4 |
| SRRT | 8 | SRRT |
| HMGCR | 9 | HMGCR |
| INSIG1 | 9 | INSIG1 |
| PGRMC1 | 9 | PGRMC1 |
| SREBF2 | 9 | SREBF2 |
| O55222 | 10 | ILK |
| P07901 | 10 | HSP90AA1 |
| P19096 | 10 | FASN |
| Q8BK64 | 10 | AHSA1 |
| BAG4 | 11 | BAG4 |
| BMX | 11 | BMX |
| CDC14A | 11 | CDC14A |
| CDC25B | 11 | CDC25B |
| CDC42 | 11 | CDC42 |
| CDK5 | 11 | CDK5 |
| CHUK | 11 | CHUK |

| | | |
|---|---|---|
| CYLD | 11 | CYLD |
| DAXX | 11 | DAXX |
| DLGAP1 | 11 | DLGAP1 |
| DYNLL1 | 11 | DYNLL1 |
| EGFR | 11 | EGFR |
| EPS15 | 11 | EPS15 |
| IKBKB | 11 | IKBKB |
| KIF20A | 11 | KIF20A |
| NCK1 | 11 | NCK1 |
| NCK2 | 11 | NCK2 |
| PAK1 | 11 | PAK1 |
| PRKCD | 11 | PRKCD |
| RAPGEF4 | 11 | RAPGEF4 |
| RASGRP3 | 11 | RASGRP3 |
| SPOPL | 11 | SPOPL |
| SPTBN2 | 11 | SPTBN2 |
| STX1A | 11 | STX1A |
| TNFRSF1A | 11 | TNFRSF1A |
| UBE2I | 11 | UBE2I |
| UNC13B | 11 | UNC13B |
| FAM98A | 12 | FAM98A |
| Q3TJZ6 | 12 | FAM98A |

***Supporting Table 5.*** *Pathway annotation results of all 13 distinct modules.*

**https://docs.google.com/spreadsheets/d/1FJSHmeTYpMRkU4UmHRfPpuJCrZybnrELAGHbwhodYNI/edit?usp=sharing**
